# A class of metallohydrolases expands bile salt hydrolase activity in the gut

**DOI:** 10.64898/2026.04.05.716488

**Authors:** Kien P. Malarney, Samantha A. Scott, Pamela V. Chang

**Author notes:** Correspondence: Prof. Pamela V. Chang, Department of Microbiology and Immunology, Department of Chemistry and Chemical Biology, 930 Campus Road, VMC C4-185, Ithaca, NY 14853.

## Abstract

Bile acids are steroidal metabolites produced by host and microbial metabolism that shape gut microbiome ecology and influence host physiology^1,2^. Bile acid structural diversification requires gut microbial bile salt hydrolase (BSH) activities, which cleave the amide bond of liver-derived bile acid amidates (BAAs)^3^. Beyond this gatekeeping metabolic function, BSHs expand the bile acid pool via their amine *N*-acyltransferase activity to produce numerous microbially derived BAAs that signal via host receptors and are further metabolized^4,5^. To date, all BSH activity has been attributed to a family of N-terminal nucleophile (Ntn) cysteine hydrolases. However, numerous gut anaerobic bacteria possess BSH activity but do not encode *bsh* genes from the Ntn family. Here, we describe a previously unknown class of <u>m</u>etal-dependent <u>BSHs</u> (mBSHs) broadly distributed in the gut microbiome. These metalloenzymes have a distinct active site architecture from the canonical Ntn superfamily of cysteine hydolase BSHs (cBSHs) and are selective for taurine-conjugated BAAs. The discovery of this heretofore unappreciated class of BSHs overturns the paradigm that this important, conserved biochemical activity of the gut microbiota is provided exclusively by canonical cysteine hydrolases, greatly expands the known landscape of bile acid metabolism, and reveals a previously unrecognized link connecting host–microbiota bile acid co-metabolism with microbial taurine utilization pathways.

## Main Text

Bile acids are major components of bile that have antimicrobial activities and mediate dietary lipid absorption in the intestines, energy metabolism, and immunity^6,7^. These amphiphilic molecules are biosynthesized in the liver from cholesterol into the primary bile acids cholic acid and chenodeoxycholic acid, which are conjugated at the C24 position to glycine and taurine to form conjugated bile acids that are stored in the gall bladder^3^. Conjugated bile acids are secreted into the small intestine to aid in digestion and are recycled to the liver in the terminal ileum via enterohepatic circulation. A small fraction of these bile acids escapes to the large intestine, where the hydroxyl groups on the steroid core are metabolized by the gut microbiota via oxidation, epimerization, and 7α-dehydroxylation, to yield numerous secondary bile acids, including deoxycholic and lithocholic acids.

Bile salt hydrolase (BSH, EC 3.5.1.24) is a cysteine hydrolase and member of the N-terminal nucleophile (Ntn) hydrolase superfamily that cleaves the amide bond within BAAs^3,8^. This enzyme carries out the initial step of secondary bile acid metabolism, which is thought to precede all subsequent bile acid biotransformations in the intestines^9,10^. Beyond this gatekeeping reaction, BSH was recently discovered to catalyze the formation of numerous microbial-derived bile acid amidates (BAAs) not limited to glyco- and tauro-conjugates via its amine *N*-acyltransferase activity (Fig. 1a), significantly broadening the diversity of BAA species and our current understanding of their metabolism in the gut^4,5^. Despite the widespread presence of BSHs in the gut microbiome, several gut bacteria, such as *Collinsella intestinalis*, *Bilophila wadsworthia*, and *Clostridioides difficile*, possess BSH activity in the absence of a known *bsh* gene^11–13^. Here, we use a chemoproteomic approach to identify a novel class of <u>m</u>etal-dependent <u>BSHs</u> (mBSHs) in the gut microbiota that, unlike canonical cysteine BSHs (cBSHs), exclusively hydrolyze taurine-conjugated bile acids to scavenge carbon and nitrogen while diversifying the overall bile acid pool within the host. These findings overturn the current dogma that BSH activity is provided exclusively by Ntn superfamily cysteine hydrolases and broaden the scope of bile acid biotransformations in the gut, which could impact metabolism, immunity, and inflammatory diseases.

**Figure 1.**
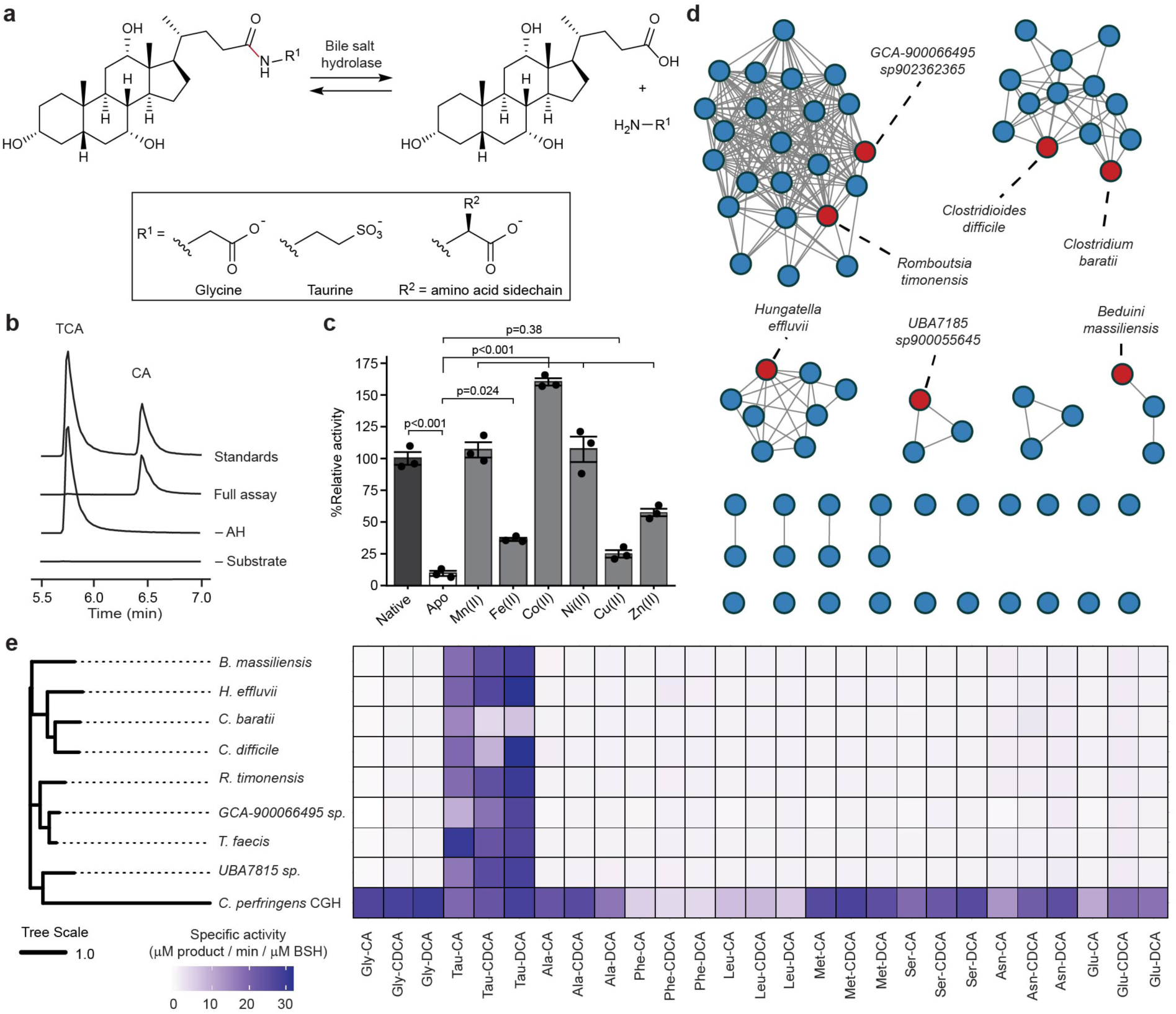
Discovery of a metal-dependent bile salt hydrolase (mBSH) from the amidohydrolase superfamily. (a) Bile salt hydrolase controls the amino acid conjugation of bile acids through conjugating and deconjugating activities. (b) Validation of BSH activity from purified *Tf* amidohydrolase by LC-MS. *Tf* amidohydrolase (0.2 mg/mL) was incubated with taurocholic acid (TCA, 5 mM) in 25 mM NaP_i_ (pH 6.8), ZnCl_2_ (100 μM), 1% (v/v) DMSO after incubation for 2.5 h at 37 °C. Extracted ion chromatograms of taurocholate (TCA) and cholate (CA) anions are shown with the same scale, with peak heights indicating relative intensities of each ion. (c) Reconstitution of *Tf* mBSH apoenzyme with metal dichloride salts rescues activity. *Tf* mBSH (20 μM) was treated with 1 mM 1,10-phenanthroline or vehicle (1% MeOH final) in 25 mM NaP_i_ (pH 6.8) for 1 h at room temperature. After buffer exchange, *Tf* mBSH apoenzyme was treated with the indicated metal dichloride in 25 mM NaP_i_ (pH 6.8, 20 μM BSH, 1 mM metal) or water. Enzyme preparations (2 μM final) were incubated with TCA (5 mM) in 25 mM NaP_i_ (pH 6.8), 1% v/v DMSO, for 1 h at 37 °C. Released taurine was quantified by ninhydrin assay. Values are reported as mean ± sem. *p*-values determined by one-way ANOVA, followed by post-hoc Tukey test. (d) Identification of *Tf* amidohydrolase homologs in the human gut microbiome. DIAMOND BLAST of *T. faecis* amidohydrolase against the Unified Human Gut Protein catalog yielded 213 proteins (*e* < 10^-150^). A sequence similarity network of the homologs was generated, with edge drawing cutoff *e* < 10^-200^ and sequences of %identity ≥ 90% collapsed into a single node. Enzymes from nodes highlighted in red were further characterized (*vide infra*). (e) *T. faecis* amidohydrolase and homologs are taurine-selective BSHs. Purified BSHs (2.5 μM) were incubated with BAA (2.5 mM) in 25 mM NaP_i_ (pH 6.8), ZnCl_2_ (100 μM), 1% (v/v) DMSO for 0.5 h at 37 °C, after which amino acid product was quantified by a ninhydrin assay. Protein phylogeny was determined using MUSCLE-derived multiple sequence alignment, followed by neighbor-joining with BLOSUM62-derived distance. BAAs are abbreviated to indicate amino acid (i.e., Tau for taurine or standard proteinogenic amino acid abbreviation) and steroid core (CA, cholic acid; CDCA, chenodeoxycholic acid; DCA, deoxycholic acid). Values reported as mean.

### *Turicibacter faecis* amidohydrolase is a metal-dependent BSH

To discover novel classes of BSHs within the gut microbiota, we used a chemoproteomic approach deploying a cholic acid-based chemical probe (**1**, Extended Data Fig. 1a) containing an electrophilic warhead and a click chemistry handle to serve as a covalent ligand for bile acid-binding proteins^14^. After labeling murine gut bacterial lysate with **1**, followed by click chemistry-based enrichment and mass spectrometry-based proteomics, *Turicibacter faecis* (*Tf*) amidohydrolase was highly enriched by the probe. Interestingly, this predicted metal-dependent hydrolase is not a member of the Ntn superfamily and is annotated as an *N*-substituted formamide deformylase (Nsfd)^15^. To assess whether this *Tf* amidohydrolase possesses noncanonical BSH activity, we first cloned, overexpressed, and purified this enzyme (Fig. S1a, Tables S1-2) and verified its labeling by **1** (Extended Data Fig. 1b,c). Enzymatic activity assays with purified *Tf* amidohydrolase revealed efficient hydrolysis of taurocholic acid (TCA) into cholic acid and taurine (Fig. 1b). Although *Tf* amidohydrolase had high homology to an annotated Nsfd, we found no evidence of hydrolysis of the cognate Nsfd substrate, *N*-benzylformamide (Extended Data Fig. 2a), suggesting that *Tf* amidohydrolase possesses bona fide BSH activity^15^. Moreover, this enzyme displayed activity in a variety of buffers over a variety of pH range and temperature (Extended Data Fig. 2b-d).

To verify the predicted metal dependence of this amidohydrolase superfamily member, we assayed BSH activity after chelator pretreatment and found its activity is sensitive to 1,10-phenanthroline treatment (Extended Data Fig. 2e). We then used 1,10-phenanthroline to generate an inactive apoenzyme whose BSH activity was rescued upon reconstitution with first-row transition metal species (Fig. 1c)^16^. Putative metal binding residues were identified based on homology to *Arthobacter pascens* Nsfd and the structure of an uncharacterized amidohydrolase from *Pyrococcus furiosus* in the Protein Databank PDB:3ICJ (Extended Data Fig. 3)^15^. On the basis of its biochemical activity as a BSH and its divalent metal cofactor dependence, we have designated this enzyme as a founding member of a new class of non-canonical BSHs termed metal-dependent BSHs (mBSHs) that are distinct from previously described cBSHs.

We identified *Tf* amidohydrolase homologs within the human gut microbiome and generated a sequence similarity network containing 212 homologs (Fig. 1d, Table S3)^17,18^. We cloned, overexpressed, and purified representative members of this family, from *Beduini massiliensis*, *Hungatella effluvii*, *Clostridium baratii*, *Clostridioides difficile*, *Romboutsia timonensis*, *GCA-900066495* sp., and *UBA7815* sp., that are at least 40% identical to *Tf* amidohydrolase based on amino acid sequence (Fig. 1e, Fig. S1b-h, Tables S1-3). We determined that, whereas a representative cBSH *Clostridium perfringens* choloylglycine hydrolase (Fig. S2a) deconjugates numerous BAAs with disparate physicochemical properties, these amidohydrolases all selectively hydrolyze taurine conjugates exclusively (Fig. 1e).

### Taurine selectivity of mBSHs depends on conserved amino acid residues

To further characterize the taurine selectivity of this novel class of mBSHs and structure-activity relationship of their preferred substrates, we tested these representative amidohydrolases with additional potential substrates, including β-alanocholic acid, an extended version of glycocholic acid with a similar length to taurocholic acid, and *N*-acetyltaurine, which shares an *N*-acyl taurine group with taurocholic acid (Fig. 2a). We found that these mBSHs act uniquely on taurine-BAAs, with preference for the sulfonate compared to other acidic groups, such as the carboxylate present in β-alanocholic acid.

**Figure 2.**
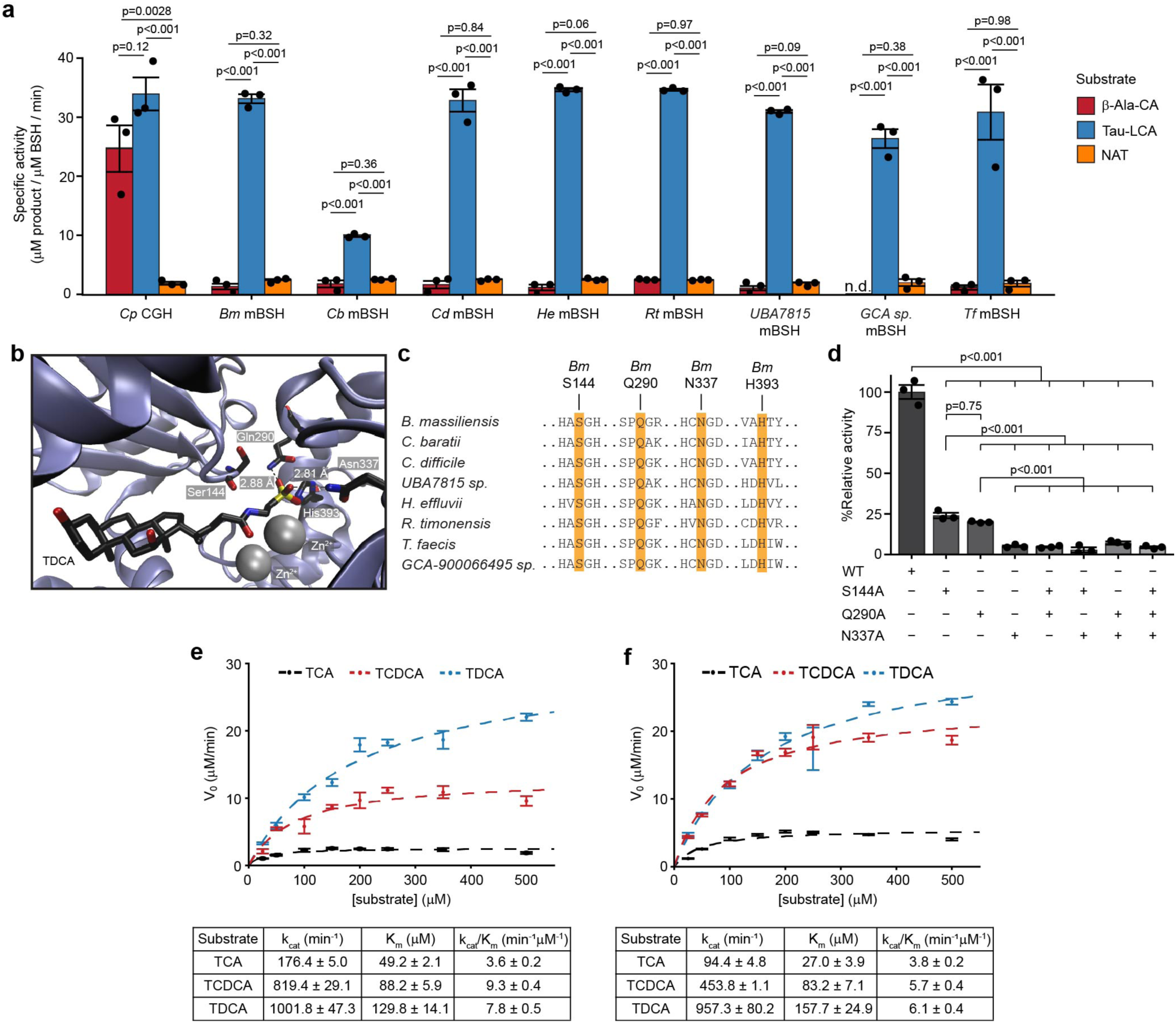
Enzymatic and structural determinants of mBSH substrate selectivity and catalysis. (a) mBSHs are selective for the sulfonate of taurine BAAs. BSHs (2.5 μM) were incubated with BAA or *N*-acetyltaurine (2.5 mM) in 25 mM NaP_i_ (pH 6.8), ZnCl_2_ (100 μM), 1% (v/v) DMSO at 37 °C for 0.5 h, after which amino acid product formation was quantified using a ninhydrin assay. Values are reported as mean ± SEM. Substrates: β-Ala-CA, β-alanocholic acid; TLCA, taurolithocholic acid; NAT, *N*-acetyltaurine. Organism abbreviations: *Cp*, *Clostridium perfringens*; *Bm, Beduini massiliensis*; *Cb*, *Clostridium baratii*; *Cd*, *Clostriodioides difficile*; *He*, *Hungatella effluvii*; *Rt*, *Romboutsia timonensis*; *UBA7815*, *UBA7815* sp900055645; *GCA sp.*, *GCA-900066495* sp902362365; *Tf*, *Turicibacter faecis*. (b) Molecular modeling of *B. massiliensis* mBSH with Zn^2+^ cofactors and substrate TDCA predicts residues involved in sulfonate recognition. Enzyme–substrate complex generated using Boltz-2. Abbreviations: TDCA, taurodeoxycholate. Atom coloring: black (carbon), red (oxygen), yellow (sulfur), blue (nitrogen), and silver (zinc). (c) Multiple sequence alignment of mBSHs identifies conservation of predicted sulfonate binding residues across characterized mBSHs. (d) Disruption of H-bonding abilities of sulfonate-binding residues attenuates *B. massiliensis* mBSH activity. Wild-type (WT) or mutant mBSH (200 nM) was incubated with TDCA (1 mM) in 25 mM NaP_i_ (pH 6.8), ZnCl_2_ (100 μM), 1% (v/v) DMSO at 37 °C for 10 min, whereafter released taurine was quantified by a ninhydrin assay. *p*-values in (a) and (d) determined by one-way ANOVA, followed by post-hoc Tukey test. (e) Kinetic characterization of *Tf* mBSH. BSH (30 nM) was incubated with taurine BAA (concentrations indicated), 25 mM NaP_i_ (pH 6.8), 100 μM ZnCl_2_, 1% (v/v) DMSO at 37 °C for 3 min. Deconjugated bile acids were quantified by LC-MS, and kinetic data were fitted to the Michaelis–Menten equation. Substrates: TCA, taurocholic acid; TCDCA, taurochenodeoxycholic acid; TDCA, taurodeoxycholic acid; TLCA, taurolithocholic acid. (f) Kinetic characterization of GCA-900066495 sp902362365 mBSH. Reaction conditions, monitoring, and data fitting were performed as in (e). Rate values in (e,f) are reported as mean ± SEM.

To understand the basis of this selectivity, we modeled *B. massiliensis* amidohydrolase complexed with taurodeoxycholic acid (TDCA) and zinc ions using Boltz-2^19^. From this structural model and multiple sequence alignment, we identified three conserved amino acid residues (Ser144, Gln290, and Asn337) involved in coordinating the sulfonate of taurine through hydrogen bonding and a conserved electrostatic interaction through His393 (Fig. 2b,c, Extended Data Fig. 4). As this enzyme exhibited selectivity for taurine-conjugated BAAs, but not β-alanocholic acid, we hypothesized that hydrogen bonding to the sulfonate group is crucial for substrate recognition. Accordingly, we performed site-directed mutagenesis of S144, Q290, and N337 to alanine (i.e., S144A, Q290A, N337A) to disrupt the interactions between these residues and the sulfonate. After overexpressing and purifying these mutant enzymes (Fig. S3), we found that TDCA deconjugation was significantly reduced compared to wild-type *B. massiliensis* amidohydrolase (Fig. 2d). We created additional double and triple alanine mutants and found that enzymatic activity toward TDCA was further reduced. These results indicate that these amino acid residues are important for conferring taurine selectivity of these mBSHs.

We also kinetically characterized mBSH from *T. faecis* (Fig. 2e) and *GCA-900066495 sp.* (Fig. 2f, Fig. S4), because these two enzymes are 85% identical based on amino acid sequence but differ in bile acid steroid core preference. We identified varying degrees of substrate inhibition (Fig. S5), though these characterizations did not fit classical substrate inhibition models. As such, we determined parameters for the region exhibiting Michaelis-Menten kinetics and found that *Tf* amidohydrolase preference for TCA can be explained by its higher k_cat_ compared to *GCA-900066495 sp.* (Table S4).

### Taurine selectivity of mBSHs correlates with taurine metabolism genes

As the *Tf* genome contained multiple canonical cBSHs in addition to mBSH, we searched for cBSHs within the genomes harboring a *Tf* mBSH homolog (Fig. 3a). To estimate gene counts, we examined cBSH and mBSH copy number in genomes with an estimated completeness ≥90%. Among these 112 genomes with an identified mBSH, approximately half lacked a cBSH, and the remaining genomes showed varying copy numbers with mBSH copy number being generally one per genome (Fig. 3b).

**Figure 3.**
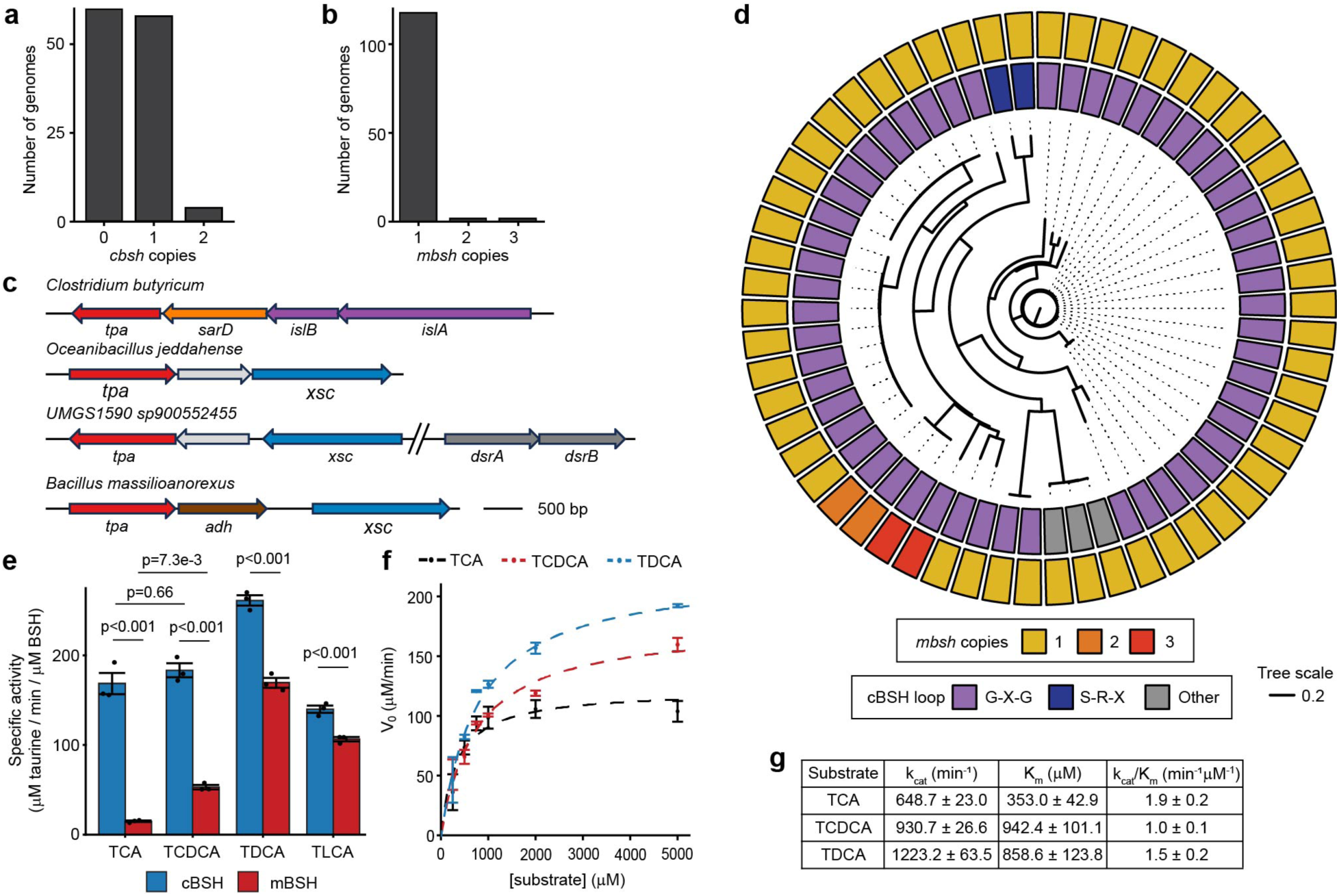
Co-occurrence of mBSH and cBSH genes reveals functional specialization within bacterial genomes. (a) Copy numbers of *cbsh* from genomes encoding an *mbsh* gene. (b) *mbsh* genes carried in these genomes. (c) Gene clusters for taurine metabolism from genomes of taxa with *mbsh*. (d) Intragenomic cBSH selectivity for genomes encoding *mbsh*. Outer ring, *mbsh* gene copies; inner ring, *cbsh* selectivity loop motif; inner tree, cBSH protein phylogeny. (e) Comparison of biochemical activities of intragenomic cBSH and mBSH from GCA-900066495 sp902362365. BSHs (200 nM) were incubated with the indicated taurine-conjugated BAA (1 mM) in 25 mM NaP_i_ (pH 6.8), 1% (v/v) DMSO at 37 °C for 15 min; released taurine product was quantified by a ninhydrin assay. Values are reported as mean ± SEM. *p*-values determined using one-way ANOVA, followed by post-hoc Tukey test. (f-g) Kinetic characterization of GCA-900066495 sp902362365 cBSH. BSH (200 nM) was incubated with the indicated taurine-conjugated BAA (0–5000 µM) in 25 mM NaP_i_ (pH 6.8), 1% (v/v) DMSO at 37 °C for 3 min. Released taurine was quantified by a ninhydrin assay; kinetic data were fitted to the Michaelis–Menten equation; values are reported as mean ± SEM.

Given the selectivity of mBSHs for taurine BAAs, we next investigated whether genomes containing *Tf* amidohydrolase homologs harbored the potential to catabolize taurine and therefore contribute to sulfur, nitrogen, and carbon metabolism (Extended Data Fig. 5, Table S5). Within these 206 genomes, corresponding to 60 different lineages, we used BLAST to identify genes involved in taurine degradation (Table S6)^20^. Of these genomes, 141 (69%) contained both Tpa and SarD homologs — catalyzing the transamination of taurine and carbonyl reduction of sulfoacetaldehyde, respectively—suggesting a potential role for mBSH aiding in nitrogen acquisition^21,22^. We also found homologs of proteins involved in carbon acquisition through pathways involving C–S lyases IslA and Xsc in 10 and 6 genomes, respectively^21,23^. These include at least four unique gene clusters involved in sulfur metabolism within diverse bacteria, including *Clostridium butyricum*, *Oceanibacillus jeddahense*, *UMGS1590 sp900552455*, and *Bacillus massilioanorexus* (Fig. 3c)^24^. As IslA and Xsc generate sulfite, an electron acceptor for anaerobic respiration, we also searched genomes for dissimilatory sulfite reductase proteins, which we found in 9 of the genomes harboring IslA or Xsc pathways^21,23,25^. Altogether, these findings suggest that taurine conjugate-selective mBSHs in human gut bacteria may facilitate dissimilatory sulfite metabolism and downstream energy harvest from taurine via nitrogen and carbon acquisition.

Interestingly, many mBSH genes are found within gut bacterial genomes that contain taurine-selective cBSHs (Fig. 3d, Table S7)^26^. To understand the metabolic advantage to bacteria of possessing both mBSHs and cBSH, we compared activities of both mBSH and cBSH from *GCA-900066495 sp.* toward a panel of taurine BAAs and found differences based on steroid core identity (Fig. 3e). Kinetic characterization of *GCA-900066495 sp.* cBSH (Fig. 3f-g, Fig. S6) revealed that these intragenomic BSHs differ in their bile acid steroid core preference (Table S4). Further, the mBSH from *GCA-900066495 sp.* was generally more catalytically efficient than the cBSH from *GCA-900066495 sp.* (Table S4). These results suggest that gut bacteria may possess an mBSH and a cBSH to provide complementarity in substrate preference for bile acid conjugates. These synergistic activities increase the overall substrate scope for deconjugation, which could confer a competitive advantage for these microbes due to the antimicrobial activities of BAAs compared to the free acid counterparts^7^.

### Domain level analysis reveals that mBSHs are widespread in the human gut microbiome

Finally, we performed sequence similarity network (SSN) analysis at the domain level for the amidohydrolase domain (Pfam: amidohydrolase 3) present in *Tf* amidohydrolase (Fig. 4a). Briefly, we used previously curated InterPro domain annotations in the Unified Human Gut Protein catalog clustered at 90% identity^17^ and assigned functions to these representative sequences based on homology to domain members with known functions (Fig. 4a, Table S8). Although our initial characterization found enzymes belonging to a multi-phylum mBSH cluster, with enzymes belonging to the phyla of *Actinobacteria*, *Bacillota*, and *Spirochaetota*, this expanded SSN analysis identified additional clusters with homology to mBSHs in other phyla, such as *Thermodesulfobacteriota* (Fig. 4a). Further analysis identified that these mBSH homologs are found in sulfite-reducing species (Fig. 4b) and have predicted selectivity for taurine-conjugated bile acids (Fig. 4c, Extended Data Fig. 6)^21^. Accordingly, we cloned, overexpressed, and purified the mBSH homolog from *Bilophila wadsworthia* (Fig. S7) and verified its selectivity for taurine-conjugated bile acids (Fig. 4d). Altogether, this domain-level SSN analysis identified the expanded distribution of 401 representative mBSHs, clustered to 90% identity, across several bacterial phyla (Fig. 4e, Table S9).

**Figure 4.**
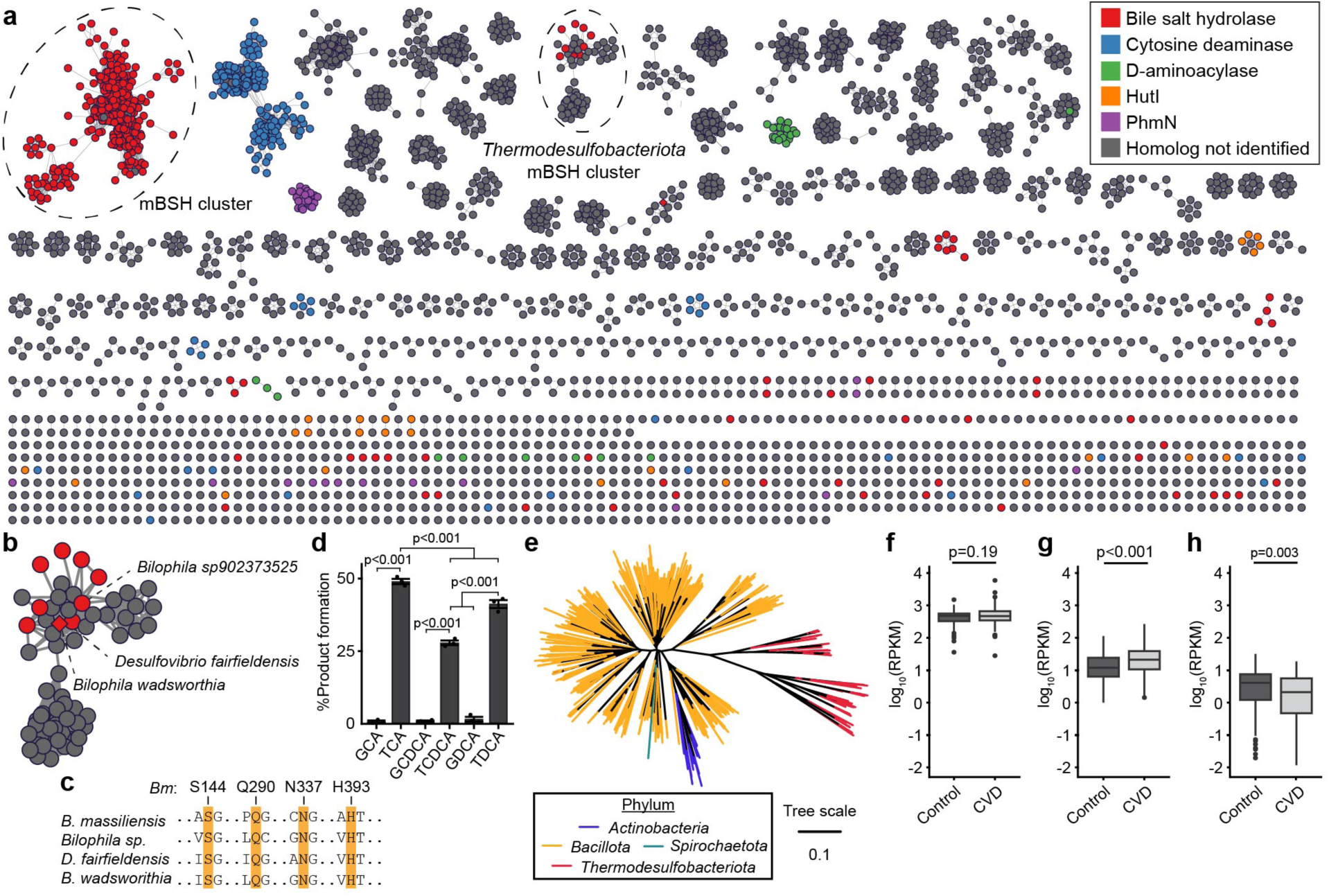
mBSH enzymes are widespread across the human gut microbiome and have increased gene abundance in cardiovascular disease (CVD). (a) Analysis of amidohydrolase 3 domain in the human gut microbiome identifies additional mBSH homologs. Proteins clustered to 90% identity from the Unified Human Gut Protein catalog containing the amidohydrolase 3 domain. A SSN of domain family members, with edge drawing cutoff *e* < 10^-150^ and sequences of %identity ≥ 90% collapsed into a single node. Protein nodes were colored according to homology with known members of the amidohydrolase 3 domain (*e* < 10^-50^, %identity ≥ 40%). Function was not assigned for proteins with high homology to enzymes that catalyze different reactions. (b) Uncharacterized *Thermodesulfobacteriota* amidohydrolase cluster identifies mBSH homologs in sulfite-reducing proteobacteria. Nodes containing proteins with high homology to mBSHs are highlighted red. (c) Multiple sequence alignment predicts *Desulfovibrionia* amidohydrolases acting on taurine conjugates. Highlighted amidohydrolases from (b) were aligned to *B. massiliensis* (*Bm*) mBSH using MUSCLE and analyzed for conserved residues involved in taurine recognition (yellow highlight). (d) *Bilophila wadsworthia* amidohydrolase is a bile salt hydrolase. Purified BSH (2.5 μM) was incubated with the indicated BAA (1 mM) in 25 mM NaP_i_ (pH=6.8) for 1 h at 37 °C. Released taurine was quantified by a ninhydrin assay. Values reported as mean ± SEM. Data are representative of n=3 independent experiments. (e) Phylogenetic tree of mBSHs from validated amidohydrolase clusters. Protein sequences from validated MBSH clusters in (a) were aligned with MUSCLE, whereafter a neighbor-joining tree was generated using distances derived from BLOSUM62. Quantitation of (f) *cbsh*, (g), multi-phylum *mbsh* cluster, and (h) *Thermodesulfobacteriota mbsh* gene family abundance in human fecal metagenomes of healthy controls (control) and atherosclerotic cardiovascular disease (CVD) patients. Genes for *cbsh* and multi-phylum *mbsh* cluster were detected in n=171 and n=214 control and CVD samples, respectively. Genes for *Thermodesulfobacteriota mbsh* cluster were detected in n=158 and n=195 control and CVD samples, respectively. p-values were determined by two-sided Mann-Whitney tests for each gene family.

The distinct taxonomic distribution and biochemical properties of mBSHs, compared to cBSHs, suggest not only functional impacts on bile acid metabolism but also associations with diseases that feature dysfunction in bile acids, taurine, and related metabolic pathways. To determine if mBSH is associated with disease, we constructed gene catalogs for representative *cbsh* and *mbsh* genes with predicted BSH activity (Tables S9-10) and quantified *bsh* gene abundances in fecal metagenomes from an atherosclerotic cardiovascular disease (CVD) study because taurine has been implicated in protection against atherosclerosis^27,28^. This *bsh* gene family-level abundance analysis identified that although *cbsh* family-level abundances did not change during CVD, *mbsh* gene abundance increased in the multi-phylum *mbsh* cluster and decreased in the *Thermodesulfobacteriota* cluster (Fig. 4f-h). Additionally, this analysis suggests that *mbsh* genes are widespread in metagenomes, albeit at a lower abundance than *cbsh* genes.

We then examined the five most widely detected *mbsh* genes to determine taxa with disease-associated *mbsh* genes for each protein cluster. Within the multi-phylum *mbsh* cluster, we identified *mbsh* genes from *Mediterranibacter gnavus* (formerly *Ruminococcus gnavus*), *Enterocloster bolteae*, *Ruthenibacterium lactatiformans*, and *Hungatella sp005845265*, whose abundances increased with CVD (Extended Data Fig. S7a-e). Meanwhile, the five most widely detected *mbsh* genes from the *Thermodesulfobacteriota* cluster were from *B. wadsworthia*, and four of these had decreased abundances in CVD patients compared to controls (Extended Data Fig. S7f-i, Table S11). These results are consistent with increased *M. gnavus* abundance and genus-level decreases in *Bilophila* abundances in CVD patients^27^.

As *M. gnavus* has previously been shown to have increased growth with supplemented taurine^29^, we searched genomes containing these CVD-associated *mbsh* genes for genes associated with taurine metabolism. BLASTP analysis identified that *M. gnavus*, *E. bolteae*, and *Hungatella sp005845265* genomes possess homologs of Tpa and SarD, suggesting the potential for these organisms to acquire nitrogen from taurine, though we did not identify any such proteins in *R. lactatiformans* genomes (Table S12, Extended Data Fig. 8). Because *Hungatella hathewayi* colonization has been shown to increase serum taurine in dysbiotic mice, the increased abundance of *Hungatella sp005845265 mbsh* may similarly modulate taurine levels in the dysbiotic CVD microbiome^30^. In summary, analysis of *bsh* gene abundance in human fecal metagenomes identified the widespread prevalence of *mbsh* genes as well as positive associations of *Clostridia* class *mbsh* genes and negative associations with *Desulfovibrionia* class *mbsh* genes with CVD that can be attributed to taurine metabolism.

## Discussion

We have identified a novel class of metal-dependent BSHs within gut microbiota that selectively catalyze the hydrolysis of taurine-conjugated bile acids. Protein domain level analysis of these amidohydrolases identified mBSH homologs across several bacterial phyla, including *Actinobacteria*, *Bacillota*, *Desulfobacterota*, and *Spirochaetota*, that are widespread in human fecal metagenomes. These results account for the longstanding, unexplainable evidence of BSH deconjugating activity in certain human gut bacteria that lack genes encoding the canonical cBSHs of the Ntn superfamily. Moreover, our bioinformatic analysis suggests that these taurine conjugate-selective mBSHs may play a role in taurine catabolism by the gut microbiota, contributing to sulfite dissimilation, nitrogen acquisition, and energy harvest for these bacteria. Ultimately, these findings highlight the widespread and functionally distinct activities of different classes of BSHs within the human gut microbiome, which broadens the scope of bile acid metabolism in the intestines.

## Methods

### Reaction Monitoring by Ninhydrin Assay

Amino acid quantitation using a ninhydrin assay was performed according to a literature procedure with minor modifications^31^. Endpoint reactions (50 μL) were quenched with 1 volume of 20% w/v trichloroacetic acid, mixed thoroughly, and pelleted by centrifugation (2207 *g*, 10 min, 4 °C). During this time, a solution of ninhydrin in 2-methoxyethanol (40 mg/mL) was prepared, which was combined with an equivalent volume of SnCl_2_ (1.6 mg/mL) in 0.2 M citrate buffer (pH = 5) to create the ninhydrin development solution.

A portion of the resulting supernatant (30 μL) was then added to 0.15 mL of ninhydrin development solution in a PCR tube or plate. The plate was then sealed and developed by heating to 95 °C for 20 min, followed by cooling for 10 min at 4 °C in a thermal cycler (Life Technologies, Waltham, MA). A portion of the developed mixture (50 μL) was then combined with water (0.15 mL) in a clear 96-well plate, and absorbance was measured at 570 nm in a BioTek Powerwave XS2 plate reader (Winooski, VT, USA). Absorbance values were corrected by absorbance values from control samples containing no enzyme. cBSH kinetic readings were corrected by subtracting absorbance values from reactions containing no substrate.

Amino acid standard solutions were prepared in the corresponding reaction buffer, quenched with 1 volume of 20% w/v trichloroacetic acid prior to analysis by ninhydrin. For cross-plate comparisons within an experiment, whole-plate absorbance values were normalized to standard solutions included in each plate so that standard absorbances were consistent across plates. Continuous monitoring was performed analogously with the modification that analysis was performed on 40 μL aliquots of reaction mixture.

### Reaction Monitoring by LC-MS

#### Mass spectrometer settings

Mass spectra were acquired with negative ion polarity with *m*/*z* range 110–1050 at 1 spectra/sec. Instrument parameters: Gas temperature, 200 °C; gas flow, 12 L/min; nebulizer, 35 psig; sheath gas temperature, 300 °C; sheath gas flow, 12 L/min. Ion source parameters: VCap, 4000 V; nozzle voltage, 1000 V; fragmentor voltage, 300 V; skimmer 1 voltage, 65 V; Octopole RF Peak, 750 V.

#### LC Method 1

Solvent A: H_2_O + 0.1% formic acid; solvent B: MeCN + 0.1% formic acid. The column was maintained at 30 °C, with a 0.5 mL/min flow rate. Compounds were eluted with the following gradient: 5% B isocratic, 0 – 0.5 min; 5 – 10% B, 0.5 – 2.20 min; 10 – 15% B, 2.20 – 2.80 min; 15 – 20% B, 2.80 – 3.10 min; 20 – 95% B, 3.10 – 7.60 min; 95 – 100% B, 7.60 – 8.00 min; 100% B isocratic, 8.00 – 10.00 min. Post-run equilibration was performed for 4 min with 5% B isocratic elution. The first 4 min of the run were directed to waste.

#### LC Method 2

Solvent A: H_2_O, 5 mM ammonium formate, 0.1% formic acid; solvent B: MeCN:H_2_O, 95:5, 5 mM ammonium formate, 0.1% formic acid. The column was maintained at 30 °C, with a 0.55 mL/min flow rate. Compounds were eluted with the following gradient: 10 – 15% B, 0.00 – 0.75 min; 15 – 45% B, 0.75 – 2.00 min; 45 – 100% B, 2.00 – 5.00 min; 100% isocratic, 5.00 – 6.75 min. Post-run equilibration was performed for 3.25 min with 10% B isocratic elution. The first 1 min of the run was directed to waste.

Bile acid species were quantified by integrating the EIC of the appropriate [M-H]^-^ ion (10 ppm mass accuracy cutoff). Bile acid molar quantities were calculated from 7-point standard curves of commercial standards.

### Cloning

Genes were codon optimized and synthesized from synthetic gene fragments (gBlocks, Integrated DNA Technologies, San Diego, CA, USA). Amplification of *mbsh* genes using a forward primer that introduced a NdeI restriction digestion site and reverse primer that introduced a C-terminal FLAG tag and XhoI restriction digestion site. The resulting amplicons were cloned into a pET21b plasmid containing an ampicillin resistance marker.

Synthetic gene fragments for *Bilophila wadsworthia mbsh* and *GCA-900066495 sp902362365 cbsh* were amplified with a forward primer that introduced a NdeI restriction digestion site and a reverse primer that introduced a XhoI restriction digestion site. The amplicons were then cloned into a pET21b plasmid harboring ampicillin resistance.

Plasmids generated during this study are indicated in Table S1. Primers used for this study are indicated in Table S2.

### Site-directed Mutagenesis

Site-directed mutagenesis was performed using the QuikChange kit (Agilent Technologies, Santa Clara, CA, USA). All mutations were performed on *Beduini massiliensis mbsh* plasmids using mutagenic primers (Table S2).

### Overexpression and Purification of Metal-dependent Bile Salt Hydrolases

Cultures of *Escherichia coli* Rosetta 2 (DE3) pLysS containing a *mbsh* plasmid were inoculated from a single colony into L-broth containing chloramphenicol (50 mg/L) and ampicillin (100 mg/L), and the cultures were grown overnight at 37 °C. The following morning, the culture was diluted into fresh L-broth containing ampicillin (100 mg/L), chloramphenicol (50 mg/L), and 100 μM ZnCl_2_ (2% subculture). The subculture was then grown to OD_600_ 0.5–0.6, at which point the culture was then chilled on ice for 20 min with periodic swirling. The culture was then treated with 0.5 M IPTG (500 μM final) and grown with shaking at 18 °C for 18-20 h.

Bacteria were then harvested by centrifugation (16,000 *g*, 4 °C, 10 min) and resuspended in ice-cold 1X PBS (pH = 7.4). The cells were pelleted once more (16,000 *g*, 4 °C, 10 min), and the cell pellets were then transferred to a 50 mL conical tube and weighed. The cells were then resuspended (9.45 mL buffer per 1 g wet pellet) in lysis buffer (20 mM Tris, 0.3 M NaCl, 20 mM imidazole, 5% v/v glycerol, 1 mM PMSF, pH = 7.4) and lysed by sonication on ice (10 s on, 20 s off, 20% amplitude, 72 cycles). The lysate was then clarified by centrifugation (16,000 *g*, 4 °C, 30 min), and the soluble fraction was separated and loaded onto a Ni-NTA column (0.7 mL slurry per 1 g wet pellet). In a 4 °C room, the lysate was then allowed to drip through the resin at a rate of 1 drop/s, after which the resin was washed with 10 bed volumes of wash buffer 1 (20 mM Tris, 0.3 M NaCl, 20 mM imidazole, 5% v/v glycerol, pH = 7.4), followed by 10 bed volumes of wash buffer 2 (20 mM Tris, 0.3 M NaCl, 50 mM imidazole, 5% v/v glycerol, pH = 7.4). Protein was then eluted with 20 mM Tris, 0.3 M NaCl, 5% v/v glycerol (pH = 7.4) buffer containing increasing concentrations of imidazole (100 mM, 150 mM, 200 mM, and 250 mM), with elution volumes of 3 bed volumes each.

The fractions were then stored at 4 °C overnight, and purity was analyzed by SDS-PAGE, followed by staining with Coomassie blue and overnight destaining. Pure fractions were then collected and concentrated using an Amicon spin column (10 kDa cutoff), which were then desalted by repeated dilution and concentration in storage buffer (20 mM Tris, 0.3 M NaCl, 5% v/v glycerol, pH = 7.4). Protein concentration was then quantified via NanoDrop using parameters calculated from ExPASY ProtParam^32^.

After quantifying total protein, the elution was aliquoted into microfuge tubes, flash-frozen with liquid nitrogen, and stored at -80 °C until further use.

### BSH Reaction Setup for Monitoring by Ninhydrin Assay

Enzymatic BSH reactions were performed in 0.2 mL PCR tubes or plates in racks or blocks prewarmed to 37 °C. Briefly, to prewarmed solutions of buffer and substrate (45 μL) was added prewarmed BSH (5 μL, 10X stock), which was then mixed thoroughly by pipetting up and down 25-30 times. Reaction tubes were then sealed and incubated at 37 °C for the specified time, after which reactions were quenched with 1 volume of 20% w/v trichloroacetic acid for analysis by ninhydrin assay. For multiple timepoint monitoring, reactions were set up analogously with larger volumes (0.24 mL), and 40 μL aliquots were quenched with 1 volume of 20% w/v trichloroacetic acid. Kinetic measurements for *GCA-900066495 sp902362365* cBSH were performed using endpoint reaction setups.

### BSH Reaction Setup for Monitoring by LC-MS

Initial characterization reactions for *T. faecis* mBSH were performed analogously to reaction setup for ninhydrin assays on a 75 μL scale, after which the reaction was quenched with 9 volumes of LC-MS grade MeOH. The quenched reaction mixture was then subjected to centrifugation (17,000 *g*, 15 min, 4 °C), and the supernatant was diluted with 49 volumes of 50% v/v MeOH, filtered using a 0.45 μm PTFE filter, and analyzed by LC-MS method A, using a 2 μL injection volume.

Initial rate determinations for *T. faecis* mBSH and for *GCA-900066495 sp902362365* mBSH were performed on a 0.24 mL scale. Kinetic reactions for *T. faecis* mBSH and for *GCA-900066495 sp902362365* mBSH were performed on a 0.1 mL scale. At each time point, reaction aliquots (40 μL) were quenched with 4 volumes of MeOH, mixed thoroughly, and stored at -20 °C for 10 min. The quenched reaction mixtures were then subjected to centrifugation (2207 *g*, 10 min, 4 °C). The supernatant was diluted with 19 volumes of 50% v/v MeOH. The diluted reaction mixtures were then subjected to centrifugation (18,000 *g*, 10 min), and the supernatant was transferred and analyzed by LC-MS. Initial rate determination and kinetic measurements were analyzed by LC-MS method B, using 1 μL and 2.5 μL injection volumes, respectively.

### Overexpression and Purification of Cysteine Hydrolase Bile Salt Hydrolases

Cultures of *Escherichia coli* Rosetta 2 (DE3) pLysS containing a cBSH plasmid were inoculated from a single colony into L-broth containing chloramphenicol and ampicillin, and the cultures were grown overnight at 37 °C. The following morning, the culture was diluted into fresh Terrific Broth containing ampicillin and chloramphenicol (2% subculture). The subculture was then grown to OD_600_ 0.5–0.6, at which point the culture was then chilled on ice for 20 min with periodic swirling. The culture was then treated with IPTG (500 μM final for *C. perfringens* cBSH and 100 μM final for *GCA-900066495 sp902362365* cBSH) and grown with shaking at 18 °C for 18-20 h.

Bacteria were then harvested by centrifugation (16,000 *g*, 4 °C, 10 min) and resuspended in ice-cold 1X PBS (pH = 7.4). The cells were pelleted once more (16,000 *g*, 4 °C, 10 min), and the cell pellets were then transferred to a 50 mL conical tube and weighed. The cells were then resuspended (9.45 mL buffer per 1 g wet pellet) in lysis buffer (1X PBS (pH = 7.4), 20 mM imidazole, 5% v/v glycerol, 1 mM PMSF, 1 mM DTT) and lysed by sonication on ice (10 s on, 20 s off, 20% amplitude, 72 cycles). The lysate was then clarified by centrifugation (16,000 *g*, 4 °C, 30 min), and the soluble fraction was separated and loaded onto a Ni-NTA column (0.7 mL slurry per 1 g wet pellet). In a 4 °C room, the lysate was then allowed to drip through the resin at a rate of 1 drop/s, after which the resin was washed with 10 bed volumes of wash buffer 1 (1X PBS (pH = 7.4), 20 mM imidazole, 5% v/v glycerol), followed by 10 bed volumes of wash buffer 2 (1X PBS (pH = 7.4), 50 mM imidazole, 5% v/v glycerol). Protein was then eluted with 1X PBS (pH = 7.4) + 5% v/v glycerol containing increasing concentrations of imidazole (100 mM, 200 mM, and 300 mM), with elution volumes of 3 bed volumes each.

The fractions were then stored at 4 °C overnight, and purity was assessed by SDS-PAGE analysis, followed by staining with Coomassie blue and overnight destaining. Pure fractions were then collected and concentrated using an Amicon spin column (10 kDa MWCO), which were then desalted by repeated dilution and concentration in storage buffer (1 X PBS (pH = 7.4) + 5% v/v glycerol). Protein concentration was then quantified by NanoDrop using parameters calculated from ExPASY ProtParam. After quantifying total protein, the elution was aliquoted into microfuge tubes, flash-frozen with liquid nitrogen, and stored at -80 °C until further use.

### Compound **1** Target Validation of *Turicibacter faecis* mBSH by In-Gel Fluorescence Imaging

In a 1.7 mL microfuge tube, 50 μg of *T. faecis* mBSH was incubated with DMSO (10% v/v final) or **1**^14^ (0.1 mM) for 1 h at 37 °C in 0.1 mL of 20 mM NaP_i_ (pH = 7) containing 1 mM DTT and 10% DMSO. The solution was treated with casein (50 μg) and subjected to chloroform – methanol precipitation, after which the protein pellet was air-dried. The resulting protein pellet was resuspended in 0.1 M NaP_i_ (pH = 7.4) + 4% SDS (94.5 μL) and subjected to copper-catalyzed azide-alkyne cycloaddition (CuAAC) using AZDye647-alkyne under previously reported conditions^14^. The protein mixture was precipitated once more, resuspended in 1X Laemmli dye containing 2-mercaptoethanol (40 μL), and resolved by SDS–PAGE. The gel was then rocked in darkness with destaining solution (AcOH:MeOH:H_2_O, 5:4:1) for 20 min, followed by rocking with water for 10 min. The gel was then imaged using a Bio-Rad ChemiDoc MP Imaging System with 647 nm excitation wavelength. The gel was then stained with Coomassie Brilliant Blue for 5 min, rinsed with water, and rocked overnight in Coomassie destaining solution (AcOH:MeOH:H_2_O 5:30:65). The gel was then imaged using a Bio-Rad ChemiDoc MP Imaging System (colorimetric mode).

Densitometry was performed with normalization to background, and fluorescence signal intensities were normalized by protein amount using Coomassie signal intensities.

### Preparation of *Turicibacter faecis* mBSH Apoenzme and Metal-Reconstituted Apoenzyme

*T. faecis* mBSH (20 μM) was incubated in 25 mM NaP_i_ (pH = 6.8) with 1 mM 1,10-phenanthroline or vehicle (1% MeOH final) at room temperature for 1 h. The enzyme was then exchanged with 50 mM NaP_i_ (pH = 6.8) using an Amicon spin column (10 kDa MWCO) by repeated dilution and concentration. Protein concentration was then quantified by the DC assay (Bio-Rad, Hercules, CA, USA).

Metal reconstitution was then performed by incubating *T. faecis* mBSH apoenzyme (20 μM) with 1 mM metal dichloride in 50 mM NaP_i_ (pH 6.8) for 10 min at room temperature. Activities of the native enzyme, apoenzyme, or metal-reconstituted apoenzyme preparations (2 μM mBSH final) were then incubated at 37 °C with 5 mM taurocholic acid in 25 mM NaP_i_ (pH = 6.8) + 1% DMSO for 1 h. Activity was assayed by quantifying released taurine by a ninhydrin assay.

## Computational Methods

### Sequence Similarity Network Generation for Homologs of *Turicibacter faecis* mBSH

The protein sequence for *Romboutsia hominis* Nsfd (MGBC100212_00368) was obtained from the MGBC^33^. This sequence was then inputted into BLASTP against the NCBI RefSeq protein database, from which the protein sequence for *Turicibacter faecis* mBSH was obtained (code: WP_161832809.1).

DIAMOND BLAST^34^ was then performed using *T. faecis* mBSH as a query for the Unified Human Gut Protein catalog^17^ clustered at 100% identity (uhgp-100), with the cutoff of *E*-value < 10^-150^. Protein sequences from the BLAST search were then obtained from uhgp-100 (Table S3) and inputted into the EFI-EST web server^35^. Sequence similarity network was generated using *E*-value < 10^-200^ for edge drawing cutoff, and sequences with 90% identity or higher were clustered into single nodes.

### Generation of Enzyme-Substrate Models using Boltz-2

Model generation was performed using Boltz-2^19^ on Colab^36^. *B. massiliensis* mBSH was modeled as a monomer using a single protein sequence. The SMILES strings for two Zn^2+^ ions ([Zn+2]) and one molecule of taurodeoxycholic acid (C[C@H](CCC(=O)NCCS(=O)(=O)O)[C@H]1CC[C@@H]2[C@@]1([C@H](C[C@H]3[C@ H]2CC[C@H]4[C@@]3(CC[C@H](C4)O)C)O)C) were used as ligand inputs. Results were visualized in VMD^37^.

### Structure-Guided Identification of cBSHs and Characterization of Selectivity Loop

A previously curated set of microbiome BSHs^26^ were aligned with MUSCLE^38^ (v5.1) and used to generate a BSH-specific HMM using HMMer^39^. This HMM was then used to search single bacterial genomes or uhgp-90 (*E*-value < 10^-10^). Sequences passing this threshold were then aligned with *C. perfringens* strain 13 BSH as a reference sequence using MUSCLE. From this alignment, candidate BSH sequences were identified based on the presence of conserved residues C2, R18, D21, N175, and R228 relative to *C. perfringens* strain 13 BSH. Identified cBSH sequences are provided in Table S7.

cBSH sequences passing this alignment-based criteria were analyzed for the selectivity loop. The identity of aligned cBSH residues relative to G211, Q212, G213 of *C. perfringens* strain 13 BSH was recorded^8^. Loops were categorized as containing ‘G-X-G’ or ‘S-R-X’ based on the identity of the aligned residues. Motifs that did not fit either category were categorized as ‘other’.

### Identification of Taurine Catabolism and Sulfidogenesis Genes in Bacterial Genomes

Protein sequences from each bacterial genome were searched for taurine catabolism and sulfidogenesis proteins using BLASTP (*e* < 10^-10^). Query sequences for IslAB pathway proteins were obtained from characterized proteins in *Bilophila wadsworthia*^21^ and *Clostridium butyricum*^24^. IslA homologs were identified as having an alignment length ≥ 800 and %identity ≥ 62% with respect to *B. wadsworthia* or *C. butyricum* IslA. The corresponding activating enzyme, IslB, was identified by homology to IslB and gene colocalization with IslA^40^. Xsc query sequences were derived from a previous survey of Xsc proteins^23^.

Tpa homologs were identified from the BLAST output by having alignment lengths of 400 amino acids or larger and %identity ≥ 28% with respect to a Tpa query. SarD homologs were identified from the BLAST output by having alignment lengths of 370 amino acids or larger and %identity ≥ 25% to a SarD query. Xsc homologs were identified from hits with alignment lengths of 550 amino acids or larger and %identity ≥ 40% to the query.

Dissimilatory sulfite reductase protein sequences, DsrA and DrsB, were obtained from *Nitratidesulfovibrio vulgaris* (strain ATCC 29579)^25^. DsrA and DsrB hits were identified by homology (40 – 60% identity) and genomic colocalization of the two proteins.

Full BLAST query sequences, names, along with their UniProt accession numbers are listed in Table S5. Resulting BLAST hits for taurine catabolism and sulfidogenesis, along with corresponding sequences and UHGG accession numbers, are provided in Table S6.

### Generation and Annotation of Amidohydrolase 3 Domain Sequence Similarity Network

Protein sequences with Interpro annotations for amidohydrolase 3 domain (pfam: PF07969) were obtained from uhgp-90 and inputted into the EFI-EST web server. Sequence similarity network was generated using *E*-value < 10^-150^ for edge drawing cutoff, and sequences with 90% identity or higher were clustered into single nodes.

Protein nodes were then annotated using BLASTP^20^. Members of the amidohydrolase 3 domain spanning various reaction types were used as query sequences. Reaction types spanned bile salt hydrolase, prolylpeptidase^41^, *N*-substituted formamide deformylase^15^, imidazolonepropionase^42^, *N*-isoproplammelide isopropyl amidohydrolase (AtzC)^43^, α-D-ribose 1-methylphosphonate 5-triphosphate pyrophosphatase^44^, cytosine deaminase^45^, 5′-deoxyadenosine deaminase^46^, and D-aminoacylases^47–50^.

BLASTP hits (*e* < 10^-50^) were sorted by reaction type. Protein reaction types were then assigned if all BLAST queries corresponded to a single reaction type and a highest %identity of alignment ≥ 40%. BLAST queries are listed by reaction type in Table S8.

### Structure-Guided Identification of mBSHs

Proteins in clusters containing mBSH homologs were aligned to *A. pascens* Nsfd^15^ using MUSCLE. Predicted active mBSHs were identified by the conservation of H69, H71, H345, H376, and D440 relative to *A. pascens* Nsfd. The corresponding sequences in multi-phylum mBSH cluster and the *Desulfovibrionia* mBSH cluster are provided in Table S9.

### Identification of Taurine-Selective mBSHs

Proteins in clusters containing mBSH homologs were aligned to *B. massiliensis* mBSH using MUSCLE. Predicted active mBSHs were identified by the conservation of S144, Q290, and N337A relative to *B. massiliensis* mBSH.

### Metagenomic and Metatranscriptomic Analysis

Metagenomic read data from Jie *et al*^27^ were obtained from the European Nucleotide Archive (PRJEB21528). Metagenomic reads were first depleted of host reads by mapping reads to the human genome (reference hg38) using Bowtie2^51^. Reads were then trimmed using trimmomatic^52^ (settings ILLUMINACLIP:TruSeq3-PE.fa:2:30:10:8:TRUE MAXINFO:80:0.5 MINLEN:50 AVGQUAL:20).

cBSH protein sequences were obtained from uhgp-90 using the previously described cBSH identified (see Structure-Guided Identification of cBSHs and Characterization of Selectivity Loop). mBSH protein sequences were obtained from the multi-phylum mBSH cluster and *Desulfovibrionia* mBSH cluster in the amidohydrolase 3 domain sequence similarity network, after which the mBSH sequences were further filtered by the presence of metal binding residues (see Structure-Guided Identification of mBSHs). Coding DNA sequences were then obtained from the corresponding genomes in the Unified Human Gut Genome catalog.

The *bsh* genes were then combined into a single catalog, and reads were then mapped to the gene catalog using bowtie2. The resulting bowtie2 output was further processed to calculate abundance as reads per kilobase million for each geneStatistical analysis for differential abundances of whole gene families or single genes were analyzed by the Mann–Whitney U test^53^ (implemented in R package ‘stats’).

To identify taurine metabolism proteins in *mbsh* genes associated with cardiovascular disease (CVD), protein cluster members for each disease-associated *mbsh* were obtained from uhgp-90. The corresponding genomes were downloaded from the Unified Human Gut Genome catalog, and the predicted protein ORFs were searched from taurine catabolism genes (see Identification of Taurine Catabolism Genes in Bacterial Genomes). BLAST hits passing criteria for homolog identification are listed in Table S10.

## Acknowledgements

We are grateful to the Arnold and Mabel Beckman Foundation (Beckman Young Investigator Award to P.V.C.) and the Alfred P. Sloan Foundation (Sloan Research Fellowship to P.V.C.) for support. This work was supported by a grant from the National Institutes of Health (NIH R35GM133501). K.P.M. was supported by an NIH Chemistry-Biology Interface Predoctoral Training Grant (T32GM138826) and a National Science Foundation GRFP (K.P.M). This work made use of the Cornell University NMR Facility, which is supported, in part, by the NSF through MRI award CHE-1531632. We thank J. Chen for technical assistance and the Weill Institute for Cell and Molecular Biology for additional resources and reagents.

## Author contributions

K.P.M. and P.V.C. conceptualized the study. K.P.M., S.A.S., and P.V.C. designed the experiments. K.P.M. synthesized the compounds, biochemically characterized the enzymes, and carried out the bioinformatic analysis. S.A.S. purified proteins and performed biochemical assays. K.P.M., S.A.S., and P.V.C. wrote the manuscript.

## Competing interests

Authors declare that they have no competing interests.

## Data availability statement

All data are available in the main text, Extended Data, and Supplementary Information. Source data are provided with this paper for Figures, Extended Data Figures, and Supplementary Figures.

## Extended Data Figures and Legends

**Extended Data Fig. 1.**
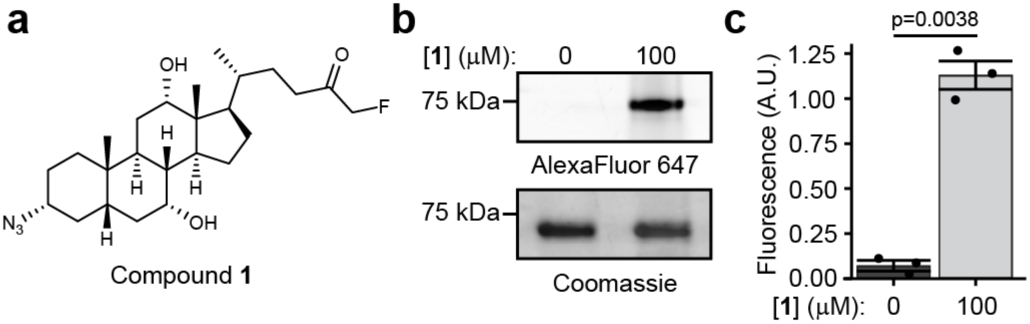
Validation of *Turicibacter faecis* mBSH labeling with compound **1**. (a) Structure of compound **1**. (b) *T. faecis* mBSH was incubated with DMSO or **1** (100 μM) for 1 h at 37 °C. The samples were tagged by CuAAC with AZDye647-alkyne and analyzed by SDS-PAGE. (c) Quantitation of fluorescence labeling after normalization by protein loading (Coomassie intensity). Values represent mean ± sem. Differences tested using 2-sided, paired *t*-test. Data representative of n=3 independent experiments.

**Extended Data Fig. 2.**
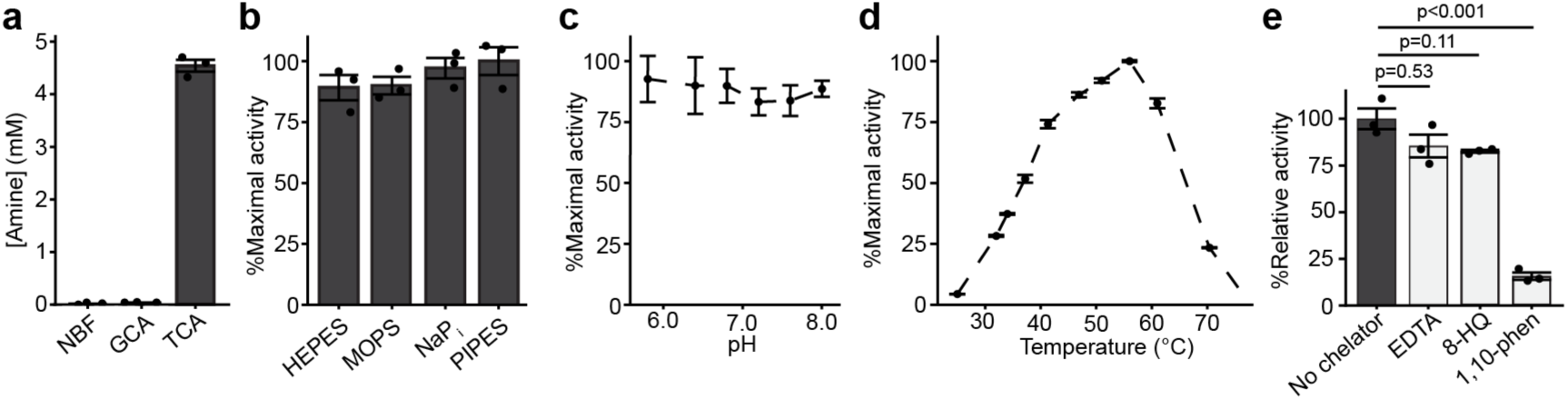
Optimization of reaction conditions for *T. faecis* T023 mBSH activity. (a) Activity after 1 h at 37 °C in NaP_i_ (25 mM, 1% DMSO, pH 6.8) with mBSH (0.2 μg/μL) and 5 mM *N*-benzylformamide (NBF), glycocholic acid (GCA), or taurocholic acid (TCA). (b) Activity after 1 h at 37 °C in different buffers (25 mM, 1% DMSO, pH = 6.8) with mBSH (0.2 μg/μL) and TCA (5 mM). (c) Activity after 1 h at 37 °C across varying pH in NaP_i_ (25 mM, 1% DMSO) with mBSH (0.2 μg/μL) and TCA (5 mM). (d) Activity in NaP_i_ (25 mM, 1% DMSO, pH = 6.8) after 15 min with mBSH (0.2 μg/μL) and TCA (5 mM). (e) Chelator dependence of *T. faecis* T023 mBSH activity. mBSH (20 μM) was incubated with chelator (1 mM) or vehicle (1% MeOH final) in 25 mM NaP_i_ (pH = 6.8) for 1 hr at room temperature, whereafter the pretreated enzyme mixtures (2 μM final) were incubated with TCA (5 mM) in 25 mM NaP_i_ (pH = 6.8, 1% DMSO) for 1 h at 37 °C. Chelator abbreviations: EDTA, ethylenediaminetetraacetic acid; 8-HQ, 8-hydroxyquinoline; 1,10-phen, 1,10-phenanthroline. Released amine in (a-e) were quantified by ninhydrin assay. *p*-values determined using one-way ANOVA, followed by post-hoc Tukey test. Data representative of *n* = 3 independent experiments. All values are reported as mean ± sem.

**Extended Data Fig. 3.**
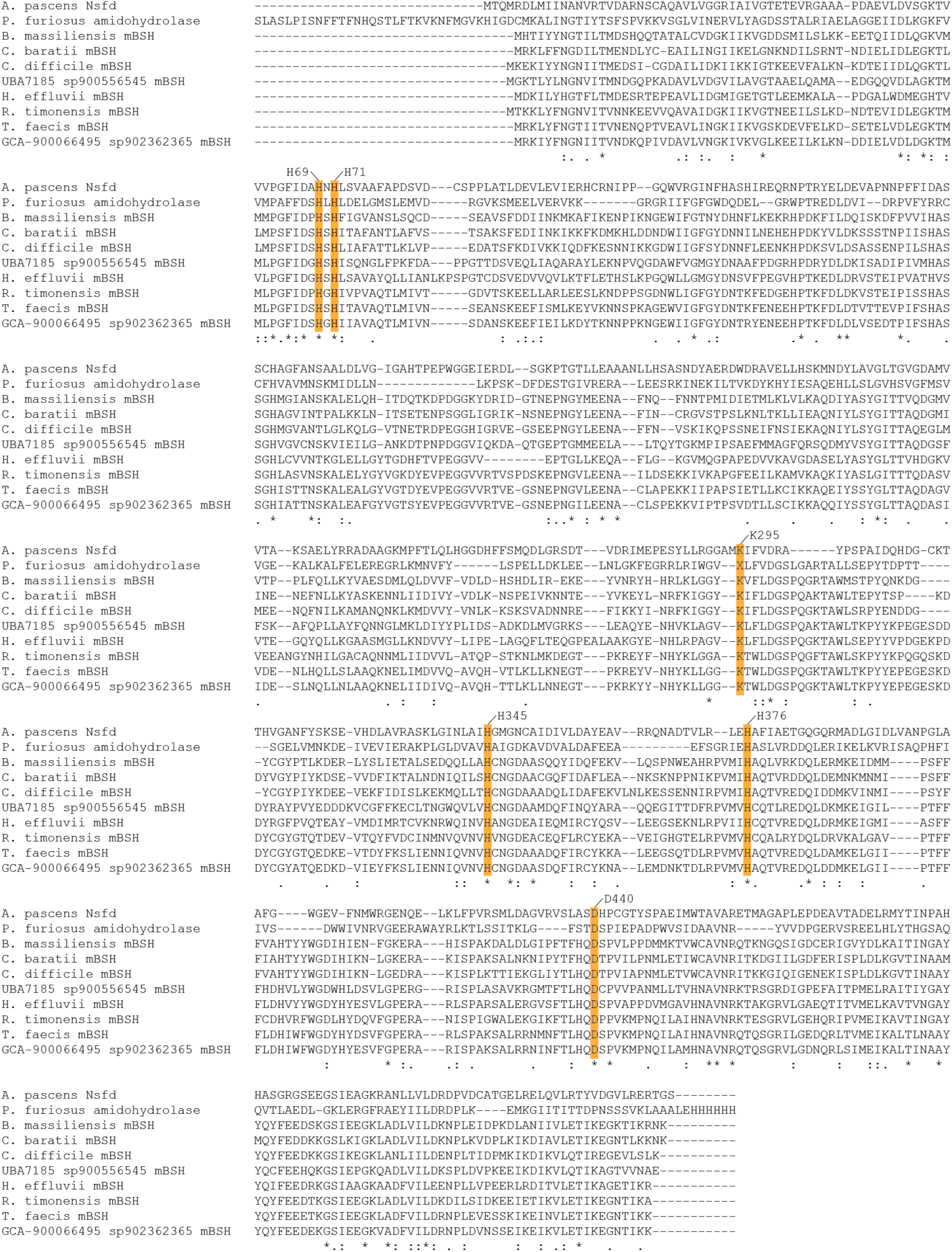
Conserved residues between *N*-substituted formamide deformylase (Nsfd), uncharacterized *P. furiosus* amidohydrolase (PDB:3ICJ), and select mBSHs. Numbered amino acid residues (yellow highlight) denote residues in *Arthrobacter pascens* Nsfd. Residue ‘X’ denotes *N*-ε-carboxy-lysine. Multiple sequence alignment was performed using MUSCLE.

**Extended Data Fig. 4.**
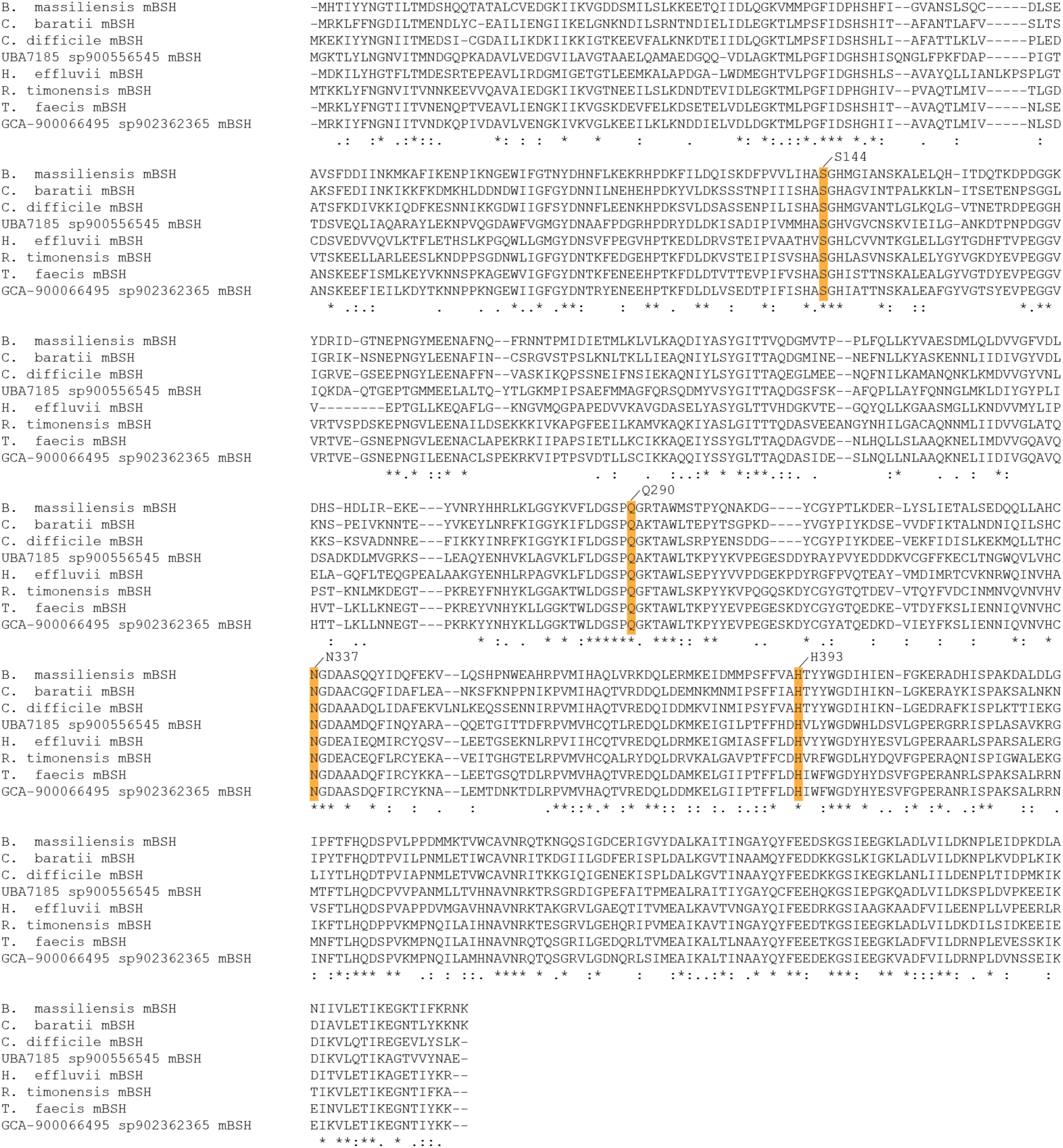
Conservation of taurine binding residues in characterized mBSHs. Numbered amino acid residues (yellow highlight) denote metal binding residues in *Beduini massiliensis* mBSH. Multiple sequence alignment was performed using MUSCLE.

**Extended Data Fig. 5.**
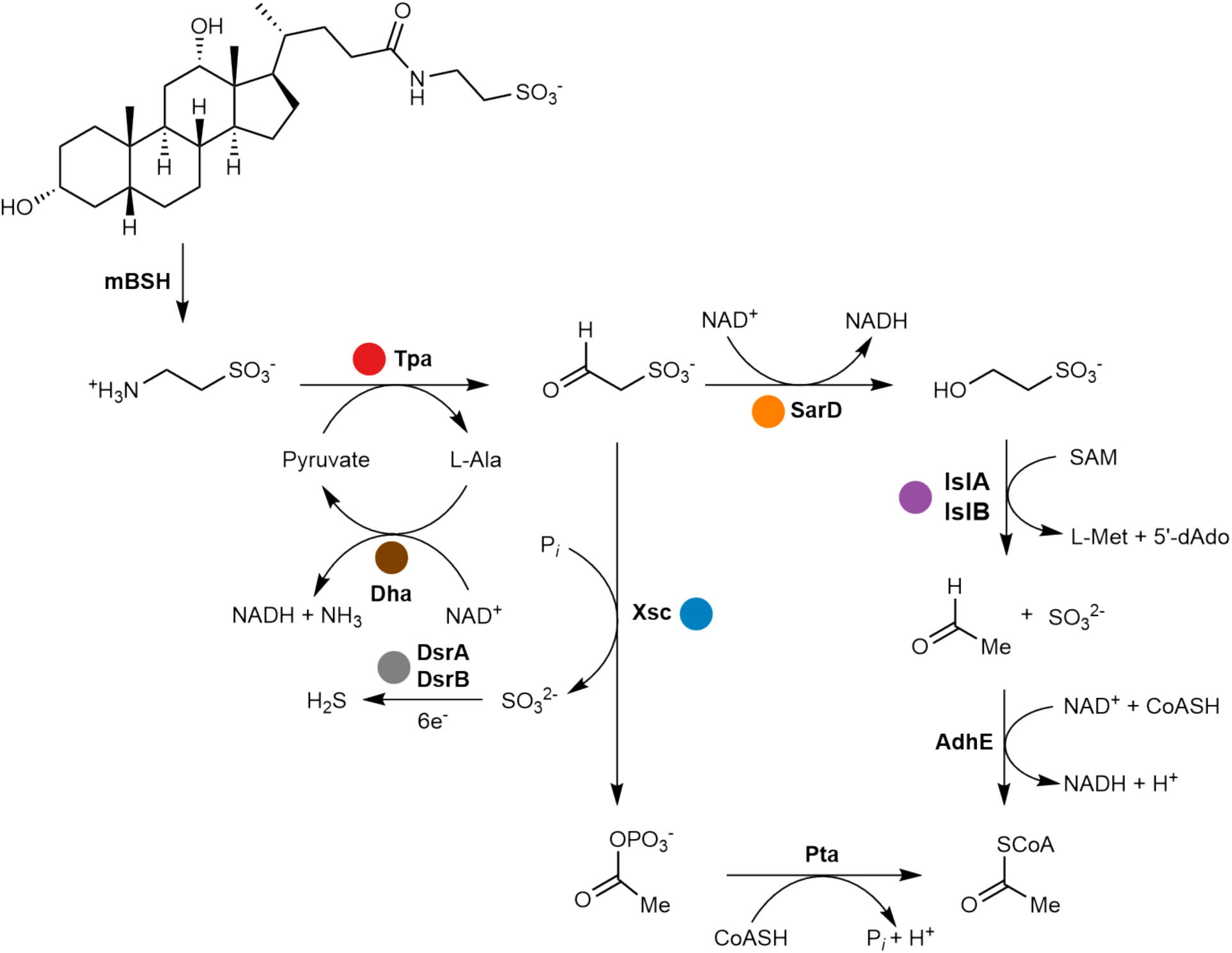
Proposed taurine assimilation pathways in bacteria with mBSH, shown from taurodeoxycholate. Protein abbreviations: mBSH, metal-dependent bile salt hydrolase; Tpa, taurine—pyruvate aminotransferase; SarD, sulfoacetaldehyde reductase; Dha, alanine dehydrogenase; Xsc, sulfoacetaldehyde acetytransferase; DsrA, dissimilatory sulfite reductase subunit α; DsrB, dissimilatory sulfite reductase submit β; IslA, isethionate—sulfite lyase; IslB, isethionate—sulfite lyase activating enzyme; AdhE, CoA-acylating acetaldehyde dehydrogenase; Pta, phosphate acetyltransferase. Chemical abbreviations: L-Ala, L-alanine; NAD^+^, nicotinamide adenine dinucleotide; NADH, reduced nicotinamide adenine dinucleotide; P*_i_*, inorganic phosphate; SAM, *S*-adenosyl-methionine; L-Met, L-methionine; 5′-dAdo, 5′-deoxyadenosine; CoA, coenzyme A.

**Extended Data Fig. 6.**
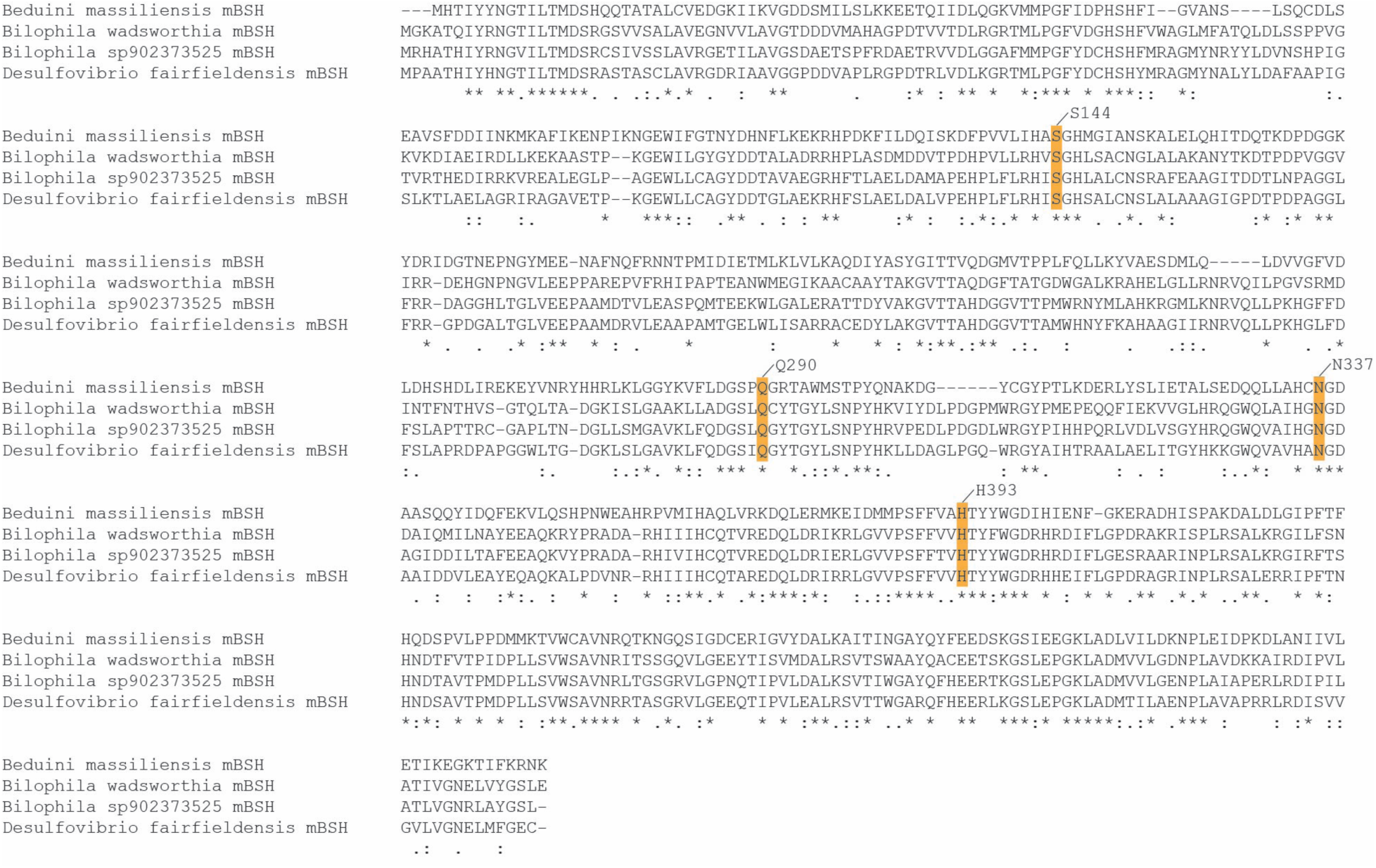
Conservation of taurine binding residues in mBSH homologs from *Desulfovibrionia* class genomes. Numbered amino acid residues (yellow highlight) denote metal binding residues in *Beduini massiliensis* mBSH. Multiple sequence alignment was performed using MUSCLE.

**Extended Data Fig. 7.**
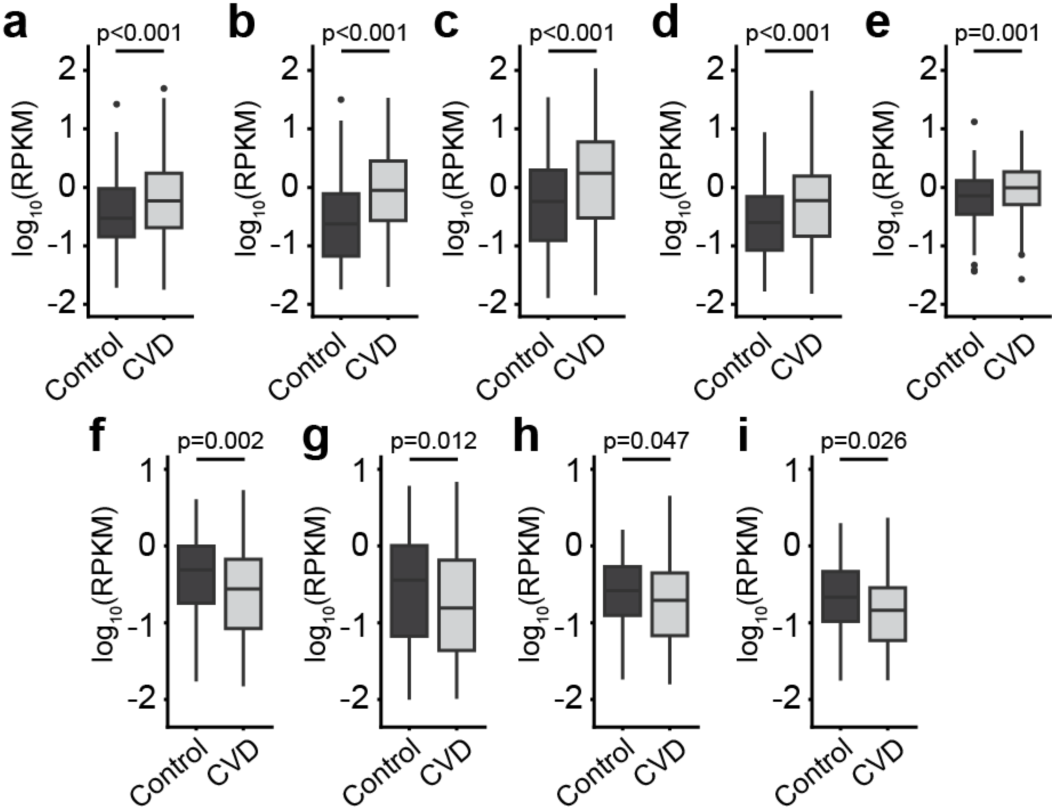
Analysis of widespread, differentially abundant *mbsh* genes in human fecal metagenomes of healthy controls (control) and atherosclerotic cardiovascular disease (CVD) patients. (a,b) Differentially abundant *mbsh* genes from *Mediterranibacter gnavus*. (c) *mbsh* gene abundance from *Ruthenibacterium lactatiformans*. (d) *mbsh* gene abundance from *Enterocloster bolteae*. (e) Differentially abundant *mbsh* gene from *Hungatella sp005845265*. (f-i) Differentially abundant *mbsh* genes from *Bilophila wadsworthia*. p-values were determined by two-sided Mann-Whitney tests for each gene.

**Extended Data Fig. 8.**
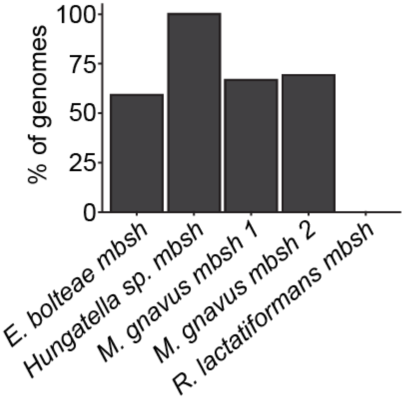
Fraction of genomes containing both Tpa and SarD homologs for genomes containing a mBSH homolog associated with cardiovascular disease. Species abbreviations: *E. bolteae*, *Enterocloster bolteae* (n=549 genomes); *Hungatella sp.*, *Hungatella sp005845265* (n=3 genomes); *M. gnavus, Mediterranibacter gnavus* (n=3 genomes for *mbsh 1*, n=1217 genomes for *mbsh 2*); *R. lactatiformans*, *Ruthenibacterium lactatiformans* (n=441 genomes).

## Notes

### Competing Interest Statement

The authors have declared no competing interest.

