## Supplementary Information for "A class of metallohydrolases expands bile salt hydrolase activity in the gut"

### Materials and Methods

#### General Considerations

Chemicals were used as received unless otherwise noted. Flash chromatography was performed using Siliaflash P60 40–63 Å 230–400 mesh silica gel obtained from SiliCycle (Quebec City, Canada). Thin-layer chromatography (TLC) was performed using glass-backed TLC 60 Å plates obtained from Merck (Burlington, MA, USA), with visualization by UV light (254 nm), followed by dipping in ceric ammonium molybdate stain and heating.

Experimental data are representative of n=3 independent experiments. Whenever possible, protein manipulations were performed on ice prior to use in enzymatic assays. Unless otherwise noted, enzymatic reactions were performed at 37 °C.

Protein sequences were obtained from the Unified Human Gut Proteins catalog (v2.0), UniProt, or NCBI. Genomes were obtained from NCBI or Unified Human Gut Genomes catalog (v2.0). Sequence similarity networks were visualized using Cytoscape[1]. Statistical analyses were performed in R (version 4.2.2) using RStudio. Kinetic data were fitted to the Michaelis – Menten equation using nonlinear least squares and are reported as the average of n=3 independent experiments. Plots were generated using cowplot and ggplot2 packages.

#### Reagent Sources

HisPur Ni-NTA resin (50% slurry), 2-methoxyethanol, imidazole, L-ascorbic acid sodium salt, and ninhydrin were obtained from Thermo Scientific (Waltham, MA, USA). IPTG was obtained from UBPBio (Dallas, TX, USA). Amicon Ultra Centrifugal filters (10 kDa MWCO) and potassium phosphate (monobasic) were obtained from Merck Millipore (Burlington, MA, USA). Sodium taurocholate hydrate, sodium phosphate (dibasic), triethylamine, and chenodeoxycholic acid were obtained from Chem-Impex (Wood Dale, IL, USA). Cholic acid was obtained from Beantown Chemical (Hudson, NH, USA). Sodium taurodeoxycholate hydrate was obtained from Spectrum Chemical (New Brunswick, NJ, USA). Sodium taurodeoxycholate hydrate was obtained from Santa Cruz Biotechnology (Dallas, TX, USA). Deoxycholic acid, glycerol, HEPES, Tris hydrochloride, sodium chloride, manganese (II) chloride tetrahydrate, sodium phosphate (monobasic), casein (sodium salt from bovine milk), trisodium citrate trihydrate, β-alanine, L-asparagine, and tetrasodium EDTA were obtained from Sigma (St. Louis, MO, USA).

*N*-benzylformamide was obtained from Ambeed (Buffalo Grove, IL, USA). Lithocholic acid and LC-MS grade ammonium formate were obtained from Sigma–Aldrich (St. Louis, MO, USA). Tin (II) chloride, zinc chloride, and MOPS were obtained from Alfa Aesar (Heysham, UK). Iron (II) chloride and ethyl chloroformate were obtained from Aldrich (St. Louis, MO, USA). PMSF was obtained from Research Products International (Mount Prospect, IL, USA). Tryptone and lysogeny broth powders were obtained from Becton Dickinson (Franklin Lakes, NJ, USA). Isopropanol, yeast extract, trichloroacetic acid, copper (II) chloride dihydrate, L-methionine, and nickel (II) chloride hexahydrate were obtained from Fisher Scientific (Pittsburgh, PA, USA). PIPES was obtained from USB (Cleveland, OH, USA). Taurine, L-glutamic acid, L-serine, L-alanine and cobalt (II) chloride hexahydrate were obtained from TCI America (Portland, OR, USA). *N*-acetyltaurine was obtained from Cayman Chemical (Ann Arbor, MI, USA). L-leucine and L-phenylalanine were obtained from Fluka (Buchs, Switzerland). Sodium dodecyl sulfate (SDS), glycine, potassium hydroxide, hydrochloric acid, and tetrahydrofuran were obtained from VWR (Radnor, PA, USA). Anhydrous sodium sulfate was obtained from EMD Chemicals (Darmstadt, Germany). Celite 545 was obtained from Acros Organics (Geel, Belgium). Sodium hydroxide was obtained from Macron Fine Chemicals (Center Valley, PA, USA). Sodium bicarbonate was obtained from Ward's Science (Rochester, NY, USA). Tris-hydroxypropyltriazolylmethylamine and AZDye 647 Alkyne were obtained from Vector Laboratories (Newark, CA, USA).

Dimethyl sulfoxide (DMSO) was obtained from Corning (Corning, NY, USA). ACS-grade acetic acid was obtained from Merck Millipore (Burlington, MA, USA). ACS-grade dichloromethane, ACS-grade chloroform, ACS-grade toluene, ACS-grade methanol, ACS-grade ethyl acetate, LC-MS grade acetonitrile, LC-MS grade formic acid, and LC-MS grade methanol were obtained from Fisher Scientific (Pittsburgh, PA, USA). Millex-LH syringe filters (0.45 µm PTFE membrane) were obtained from Millipore (Burlington, MA, USA). DMSO-*d*6 and MeOH-*d*4 were obtained from Cambridge Isotope Laboratories (Tewksbury, MA, USA).

Triethylamine was dried over potassium hydroxide, distilled under reduced pressure, and stored over potassium hydroxide pellets in a desiccator box prior to use.

Analytical Instrumentation

NMR spectra were acquired on a Bruker AVIII-500 spectrometer. $^{1}$H NMR spectra were acquired at 500 MHz, and $^{13}$C NMR spectra were acquired at 126 MHz. Chemical shifts are reported in ppm relative to the solvent residual; coupling constants *J* are reported in Hz. LC-MS analysis was performed using an Agilent 6546 LC/Q-TOF mass spectrometer coupled to an Agilent 1260 HPLC equipped with an Agilent InfinityLab Poroshell 120 EC-C18 column (3.0 mm × 50 mm, 2.7 µm mesh). Nanodrop readings were performed using a NanoDrop OneC instrument obtained from Thermo Scientific (Waltham, MA, USA).

**Chemical Synthesis**

General Procedure 1: Synthesis of Bile Acid Amidates

Conjugated bile acid synthesis was performed based on a previously describe procedure[2]. To a 50 mL round-bottom flask charged with a stir bar was added bile acid (0.48 mmol, 1.00 equiv.), followed by THF (6 mL, 0.08 M) and triethylamine (80 µL, 0.576 mmol, 1.20 equiv.). Ethyl chloroformate (80 µL, 0.576 mmol, 1.20 equiv.) was then added dropwise, and the flask was sealed with a cap. The resulting white slurry was then stirred at room temperature for 90 min.

The reaction was then treated with amino acid (0.740 mmol, 1.54 equiv.) in an aqueous solution (6 mL) containing NaOH (0.72 mmol, 1.50 equiv.) and stirred for 2 h at room temperature. The reaction was then quenched with 1 M HCl (25 mL) and partitioned with EtOAc. The aqueous layer was extracted with 2 x 20 mL EtOAc, and the combined organic layers were then washed with brine (50 mL) and dried over $Na_2SO_4$. The dried organic layers were then filtered, slurried with celite, and concentrated to dryness. The crude celite was then loaded onto a flash column. The product was then purified by flash chromatography.

Bile acid amidates of α-amino acids were synthesized using the corresponding L-amino acid enantiomer.

Characterization of Synthesized Molecules

**Choloyl alanine (Ala-CA)**

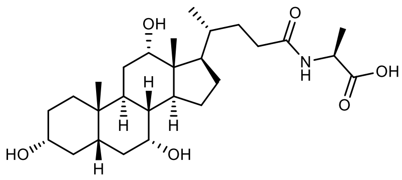

Ala-CA was synthesized according to general procedure 1 on a 0.24 mmol scale. Purification by FCC (10→12.5→15→20% MeOH/DCM + 1% AcOH) furnished a white foam (49 mg, 43% yield). Characterization was consistent with previous literature[3].

**Chenodeoxycholoyl alanine (Ala-CDCA)**

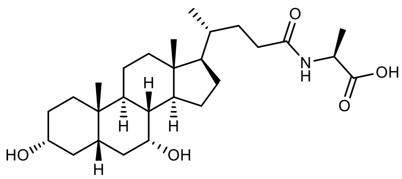

Ala-CDCA was synthesized according to general procedure 1. Purification by FCC (7→9% MeOH/DCM + 1% AcOH) furnished a white foam (165 mg, 72% yield).

Analytical data: $^{1}$H NMR (500 MHz, MeOD) δ 4.35 (q, *J* = 7.3 Hz, 1H), 3.79 (q, *J* = 2.8 Hz, 1H), 3.42 – 3.33 (m, 1H), 2.33 – 2.22 (m, 2H), 2.15 (ddd, *J* = 13.9, 9.7, 6.6 Hz, 1H), 2.05 – 1.78 (m, 6H), 1.74 (dtt, *J* = 9.1, 6.5, 2.7 Hz, 1H), 1.69 – 1.57 (m, 2H), 1.51 (tdd, *J* = 15.3, 8.0, 3.7 Hz, 5H), 1.38 (d, *J* = 7.1 Hz, 4H), 1.36 – 1.24 (m, 5H), 1.24 – 1.04 (m, 3H), 0.98 (d, *J* = 6.5 Hz, 4H), 0.93 (s, 3H), 0.70 (s, 3H). $^{13}$C NMR (126 MHz, DMSO) δ 174.46, 172.28, 70.33, 66.16, 55.60, 54.92, 50.02, 47.52, 41.91, 41.42, 35.31, 35.00, 34.83, 34.74, 32.28, 32.05, 31.47, 30.56, 27.80, 23.16, 22.72, 20.26, 18.38, 17.38, 11.66. HR-MS (ESI-TOF-MS): $[M-H]^-$ calcd. for $C_{27}H_{44}NO_5^-$ 462.3224; found 462.3230.

**Deoxycholoyl alanine (Ala-DCA)**

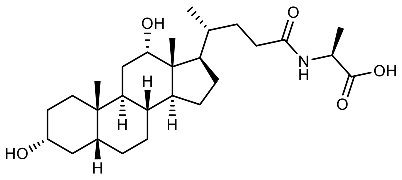

Ala-DCA was synthesized according to general procedure 1. Purification by FCC (7→9% MeOH/DCM + 1% AcOH) furnished a white foam (192 mg, 84% yield).

Analytical data: $^{1}$H NMR (500 MHz, MeOD) δ 4.36 (q, *J* = 7.3 Hz, 1H), 3.96 (t, *J* = 2.9 Hz, 1H), 3.52 (tt, *J* = 11.2, 4.6 Hz, 1H), 2.36 – 2.22 (m, 1H), 2.16 (ddd, *J* = 13.9, 9.7, 6.7 Hz, 1H), 1.97 – 1.72 (m, 7H), 1.70 – 1.55 (m, 3H), 1.55 – 1.23 (m, 14H), 1.22 – 1.05 (m, 2H), 1.03 (d, *J* = 6.5 Hz, 3H), 1.01 – 0.95 (m, 1H), 0.93 (s, 3H), 0.71 (s, 3H). $^{13}$C NMR (126 MHz, DMSO) δ 174.43, 172.43, 71.04, 69.96, 54.92, 48.61, 47.48, 47.43, 46.23, 45.99, 41.63, 36.31, 35.67, 35.16, 35.06, 33.83, 32.93, 32.15, 31.56, 30.24, 28.62, 27.22, 27.00, 26.13, 23.53, 23.11, 17.28, 17.12, 12.45. HR-MS (ESI-TOF-MS): $[M-H]^-$ calcd. for $C_{27}H_{44}NO_5^-$ 462.3224; found 462.3217.

**Choloyl phenylalanine (Phe-CA)**

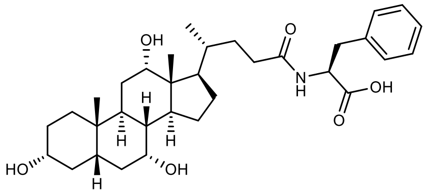

Phe-CA was synthesized according to general procedure 1. Purification by FCC (10→12→14% MeOH/DCM + 1% AcOH) furnished a white foam (155 mg, 58% yield). Characterization was consistent with previous literature[4].

**Chenodeoxycholoyl phenylalanine (Phe-CDCA)**

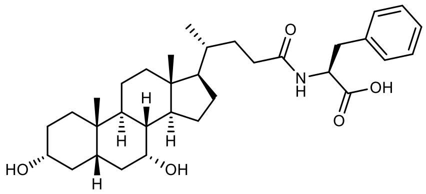

Phe-CDCA was synthesized according to general procedure 1. Purification by FCC (7→9% MeOH/DCM + 1% AcOH) furnished a white foam (125 mg, 48% yield). Characterization was consistent with previous literature[5].

**Deoxycholoyl phenylalanine (Phe-DCA)**

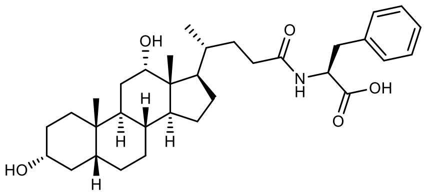

Phe-DCA was synthesized according to general procedure 1. Purification by FCC (5.5→6.5→7% MeOH/DCM + 1% AcOH) furnished a white foam (183 mg, 71% yield). Characterization was consistent with previous literature[5].

**Choloyl leucine (Leu-CA)**

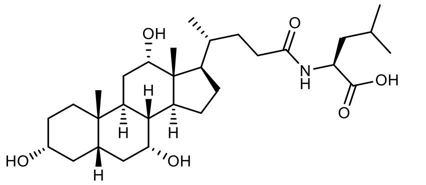

Leu-CA was synthesized according to general procedure 1. Purification by FCC (9→12→15% MeOH/DCM + 1% AcOH) furnished a white foam (166 mg, 66% yield). Characterization was consistent with previous literature[4].

**Chenodeoxycholoyl leucine (Leu-CDCA)**

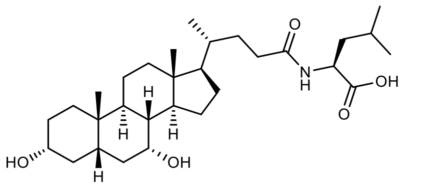

Leu-CDCA was synthesized according to general procedure 1. Purification by FCC (8→9→11% MeOH/DCM + 1% AcOH) furnished a white foam (140 mg, 58% yield). Characterization was consistent with previous literature[6].

**Deoxycholoyl leucine (Leu-DCA)**

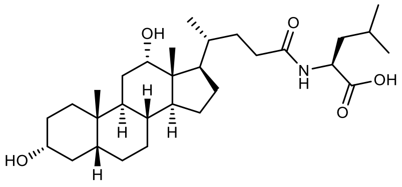


Leu-DCA was synthesized according to general procedure 1. Purification by FCC (8→9→11% MeOH/DCM + 1% AcOH) furnished a white foam (180 mg, 68% yield). Characterization was consistent with previous literature[6].

**Choloyl methionine (Met-CA)**

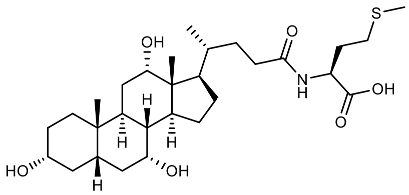


Met-CA was synthesized according to general procedure 1. Purification by FCC (7→9→10% MeOH / $CH_2Cl_2$ + 1% AcOH) furnished a white solid (65 mg, 36% yield).

Analytical data: $^1$H NMR (500 MHz, MeOD) δ 4.47 (s, 1H), 3.96 (d, $J$ = 3.0 Hz, 1H), 3.80 (q, $J$ = 3.0 Hz, 1H), 3.38 (dt, $J$ = 8.7, 4.4 Hz, 1H), 2.53 (td, $J$ = 12.2, 6.3 Hz, 2H), 2.39 – 2.22 (m, 3H), 2.22 – 2.11 (m, 2H), 2.09 (s, 3H), 2.04 – 1.70 (m, 8H), 1.70 – 1.49 (m, 6H), 1.49 – 1.26 (m, 6H), 1.12 (dd, $J$ = 12.2, 5.7 Hz, 1H), 1.04 (d, $J$ = 6.5 Hz, 3H), 0.98 (td, $J$ = 14.1, 3.4 Hz, 1H), 0.92 (s, 3H), 0.72 (s, 3H). $^{13}$C NMR (126 MHz, DMSO) δ 172.57, 71.02, 70.43, 66.24, 54.92, 51.56, 48.60, 46.22, 45.75, 35.32, 35.18, 34.89, 34.39, 32.50, 31.75, 31.27, 30.41, 29.83, 28.55, 27.35, 26.21, 22.83, 22.64, 17.13, 14.61, 12.36. HR-MS (ESI-TOF-MS): $[M-H]^-$ calcd. for $C_{29}H_{49}NO_6S^-$ 538.3207; found 538.3202.

**Chenodeoxycholoyl methionine (Met-CDCA)**

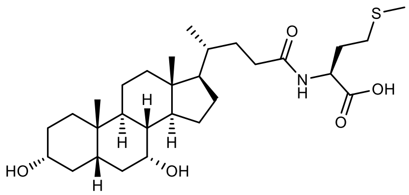


Met-CDCA was synthesized according to general procedure 1. Purification by FCC (7→8% MeOH/$CH_2Cl_2$ + 1% AcOH) furnished a white solid (133 mg, 53% yield). Characterization was consistent with previous literature[5].

**Deoxycholoyl methionine (Met-DCA)**

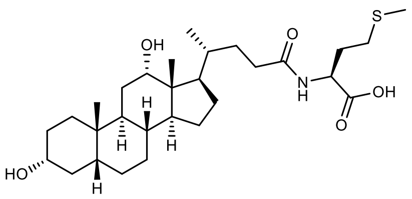

Met-DCA was synthesized according to general procedure 1. Purification by FCC (7→8→9% MeOH/$CH_2Cl_2$ + 1% AcOH) furnished a white solid (172 mg, 68% yield). Characterization was consistent with previous literature[5].

**Choloyl serine (Ser-CA)**

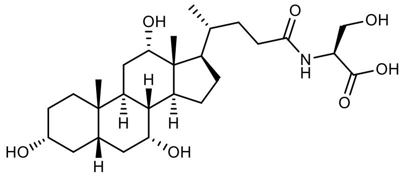

Ser-CA was synthesized according to general procedure 1. Purification by FCC (12→15→18% MeOH/$CH_2Cl_2$ + 1% AcOH) furnished a white solid (50 mg, 21% yield). Characterization was consistent with previous literature[3].

**Chenodeoxycholoyl serine (Ser-CDCA)**

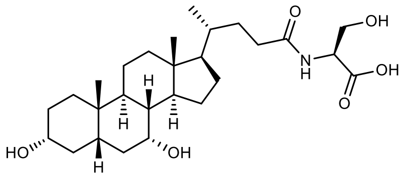

Ser-CDCA was synthesized according to general procedure 1. Purification by FCC (12→15% MeOH/$CH_2Cl_2$ + 1% AcOH) furnished a white solid (140 mg, 66% yield).

Analytical data: $^1$H NMR (500 MHz, MeOD) δ 4.43 (t, *J* = 4.6 Hz, 1H), 3.93 – 3.77 (m, 3H), 3.37 (td, *J* = 6.8, 3.4 Hz, 1H), 2.39 – 2.15 (m, 3H), 2.02 (t, *J* = 3.3 Hz, 1H), 1.99 (s, 2H), 1.98 – 1.79 (m, 5H), 1.74 (dddd, *J* = 11.7, 9.2, 5.9, 2.6 Hz, 1H), 1.70 – 1.58 (m, 2H), 1.56 – 1.43 (m, 5H), 1.41 – 1.26 (m, 6H), 1.25 – 1.05 (m, 3H), 1.04 – 0.95 (m, 4H), 0.93 (s, 3H), 0.70 (s, 3H). $^{13}$C NMR (126 MHz, DMSO) δ 172.57, 172.11, 70.34, 66.17, 61.70, 55.65, 54.92, 54.65, 50.02, 48.61, 41.93, 41.43, 35.32, 35.08, 34.84, 34.75, 32.29, 32.19, 31.50, 30.57, 27.83, 23.18, 22.73, 21.14, 20.27, 18.39, 11.68. HR-MS (ESI-TOF-MS): $[M-H]^-$ calcd. for $C_{27}H_{44}NO_7^-$ 478.3174; found 478.3176.

**Deoxycholoyl serine (Ser-DCA)**

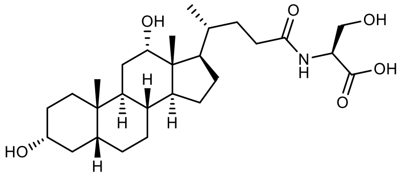


Ser-DCA was synthesized according to general procedure 1. Purification by FCC (12→15% MeOH/$CH_2Cl_2$ + 1% AcOH) furnished a white solid (140 mg, 61% yield).

Analytical data: $^1$H NMR (500 MHz, MeOD) δ 4.42 (d, *J* = 4.7 Hz, 1H), 4.00 – 3.94 (m, 1H), 3.84 (ddd, *J* = 33.2, 11.1, 4.5 Hz, 2H), 3.52 (tt, *J* = 11.1, 4.6 Hz, 1H), 2.35 (ddd, *J* = 14.9, 10.2, 5.1 Hz, 1H), 2.20 (ddd, *J* = 14.0, 9.7, 6.4 Hz, 1H), 1.97 – 1.72 (m, 7H), 1.61 (q, *J* = 7.4 Hz, 3H), 1.53 (dt, *J* = 9.1, 2.9 Hz, 2H), 1.51 – 1.34 (m, 7H), 1.34 – 1.23 (m, 2H), 1.22 – 1.07 (m, 2H), 1.04 (d, *J* = 6.4 Hz, 3H), 0.98 (td, *J* = 14.1, 3.4 Hz, 1H), 0.93 (s, 3H), 0.71 (s, 3H). $^{13}$C NMR (126 MHz, DMSO) δ 172.56, 71.03, 69.95, 61.79, 54.92, 54.67, 48.60, 47.47, 46.28, 46.00, 41.62, 36.30, 35.66, 35.15, 33.83, 32.93, 32.38, 31.58, 30.24, 28.61, 27.23, 27.00, 26.12, 23.54, 23.11, 17.13, 12.46. HR-MS (ESI-TOF-MS): $[M-H]^-$ calcd. for $C_{27}H_{44}NO_7^-$ 478.3174; found 478.3165.

**Choloyl asparagine (Asn-CA)**

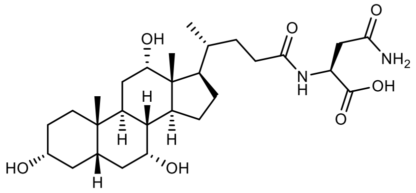


Asn-CA was synthesized according to general procedure 1. Purification by FCC (20→25% MeOH/DCM + 1% AcOH) furnished a tan foam (87 mg, 35% yield).

Analytical data: $^1$H NMR (500 MHz, MeOD) δ 4.51 (s, 1H), 3.95 (t, *J* = 2.9 Hz, 1H), 3.80 (q, *J* = 3.0 Hz, 1H), 3.38 (td, *J* = 6.7, 3.3 Hz, 1H), 2.85 – 2.64 (m, 2H), 2.39 – 2.22 (m, 3H), 2.16 (ddd, *J* = 14.7, 9.7, 6.1 Hz, 1H), 2.00 (td, *J* = 11.5, 6.8 Hz, 2H), 1.94 (s, 2H), 1.93 – 1.70 (m, 5H), 1.69 – 1.50 (m, 6H), 1.48 – 1.34 (m, 4H), 1.34 – 1.24 (m, 3H), 1.12 (tt, *J* = 11.9, 5.3 Hz, 1H), 1.03 (d, *J* = 6.3 Hz, 3H), 0.97 (dd, *J* = 14.1, 3.5 Hz, 1H), 0.92 (s, 3H), 0.71 (s, 3H). $^{13}$C NMR (126 MHz, DMSO) δ 173.14, 172.13, 71.01, 70.42, 66.24, 50.78, 48.58, 46.17, 45.75, 41.53, 41.38, 35.38, 35.33, 34.91, 34.41, 32.68, 31.63, 30.41, 28.57, 27.37, 26.23, 22.85, 22.65, 17.19, 12.39. HR-MS (ESI-TOF-MS): $[M-H]^-$ calcd. for $C_{28}H_{45}N_2O_7^-$ 521.3232; found 521.3239.

**Chenodeoxycholoyl asparagine (Asn-CDCA)**

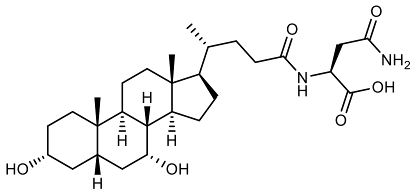

Asn-CDCA was synthesized according to general procedure 1. Purification by FCC (12→15→18% MeOH/DCM + 1% AcOH) furnished a white foam (97 mg, 40% yield).

Analytical data: $^{1}H$ NMR (500 MHz, MeOD) δ 4.65 (t, *J* = 6.1 Hz, 1H), 3.79 (q, *J* = 2.8 Hz, 1H), 3.38 (dt, *J* = 11.3, 4.3 Hz, 1H), 2.74 (qd, *J* = 15.6, 5.9 Hz, 2H), 2.35 – 2.22 (m, 2H), 2.16 (ddd, *J* = 14.0, 9.8, 6.6 Hz, 1H), 2.01 (d, *J* = 3.4 Hz, 1H), 1.99 (d, *J* = 2.8 Hz, 1H), 1.98 – 1.69 (m, 6H), 1.63 (ddt, *J* = 19.0, 12.1, 2.9 Hz, 2H), 1.55 – 1.43 (m, 5H), 1.40 – 1.26 (m, 6H), 1.24 – 1.06 (m, 3H), 1.03 – 0.95 (m, 4H), 0.93 (s, 3H), 0.69 (s, 3H). $^{13}C$ NMR (126 MHz, DMSO) δ 173.64, 172.22, 171.92, 70.33, 66.18, 55.63, 50.01, 49.44, 48.60, 41.92, 41.43, 37.65, 35.32, 35.07, 34.83, 34.75, 32.28, 31.49, 30.56, 27.82, 23.18, 22.73, 20.27, 18.37, 11.68. HR-MS (ESI-TOF-MS): $[M-H]^-$ calcd. for $C_{28}H_{45}N_2O_6^-$ 505.3283; found 505.3280.

**Deoxycholoyl asparagine (Asn-DCA)**

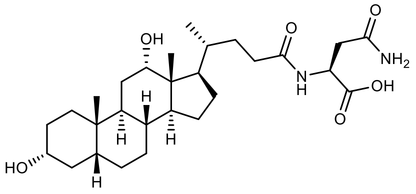

Asn-DCA was synthesized according to general procedure 1 with modifications. Briefly, after aqueous workup, the combined ethyl acetate layers were stored at 4 °C overnight, during which time the product crystallized as white needles (163 mg, 67% yield).

Analytical data: $^{1}H$ NMR (500 MHz, MeOD) δ 4.71 (dd, *J* = 6.8, 5.3 Hz, 1H), 3.96 (t, *J* = 2.9 Hz, 1H), 3.57 – 3.48 (m, 1H), 2.82 – 2.69 (m, 2H), 2.30 (ddd, *J* = 14.7, 10.0, 5.1 Hz, 1H), 2.17 (ddd, *J* = 13.9, 9.6, 6.7 Hz, 1H), 1.95 – 1.74 (m, 7H), 1.67 – 1.55 (m, 3H), 1.55 – 1.23 (m, 11H), 1.22 – 1.05 (m, 2H), 1.02 (d, *J* = 6.6 Hz, 3H), 0.97 (dd, *J* = 14.1, 3.5 Hz, 1H), 0.93 (s, 3H), 0.71 (s, 3H). $^{13}C$ NMR (126 MHz, DMSO) δ 173.04, 172.48, 171.25, 71.04, 69.95, 48.72, 48.61, 47.47, 46.26, 45.99, 41.61, 36.80, 36.31, 35.66, 35.16, 35.05, 33.83, 32.93, 32.25, 31.55, 30.24, 28.61, 27.22, 27.00, 26.12, 23.53, 23.11, 17.10, 12.47. HR-MS (ESI-TOF-MS): $[M-H]^-$ calcd. for $C_{28}H_{45}N_2O_6^-$ 505.3283; found 505.3271.

**Choloyl glutamic acid (Glu-CA)**

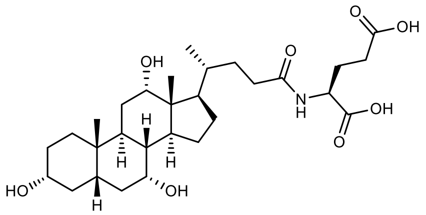

Glu-CA was synthesized according to general procedure 1. Purification by FCC (9→12→15% MeOH/DCM + 1% AcOH) furnished a white foam (46 mg, 18% yield). Characterization was consistent with previously reported literature[5].

**Chenodeoxycholoyl glutamic acid (Glu-CDCA)**

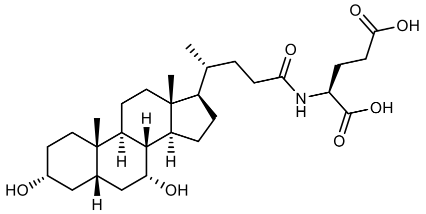

Glu-CDCA was synthesized according to general procedure 1 with modifications. Briefly, the product was allowed to crystallize from the combined ethyl acetate layers at 4 °C, and the resulting crystals were dissolved in MeOH/DCM (1:3) and filtered over celite. The filtrate was then concentrated to furnish a white powder (64 mg, 26% yield). Characterization was consistent with previously reported literature[5].

**Deoxycholoyl glutamic acid (Glu-DCA)**

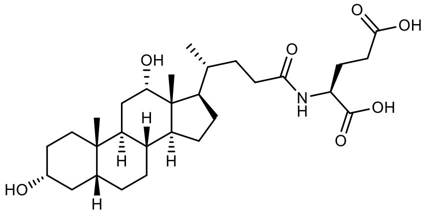

Glu-DCA was synthesized according to general procedure 1 with modifications. Briefly, the reaction was stirred for 3.25 h after the addition of amino acid. Purification by FCC (6→9→11% MeOH/DCM + 1% AcOH) furnished a white foam (49 mg, 20% yield). Characterization was consistent with previously reported literature[5].

**Choloyl β-alanine (β-Ala-CA)**

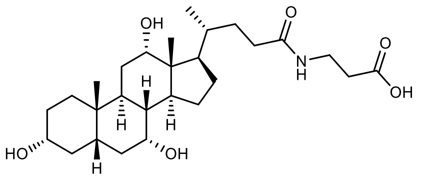

β-Ala-CA was synthesized according to general procedure 1 on a 0.24 mmol scale. Purification by FCC (6→8→9→10% MeOH/DCM + 1% AcOH) furnished a white foam (59 mg, 51% yield).

Analytical data: $^1$H NMR (500 MHz, MeOD) δ 3.95 (t, *J* = 3.0 Hz, 1H), 3.80 (q, *J* = 3.0 Hz, 1H), 3.40 (t, *J* = 6.7 Hz, 2H), 3.37 (q, *J* = 3.2 Hz, 1H), 2.47 (t, *J* = 6.7 Hz, 2H), 2.34 – 2.20 (m, 3H), 2.10 (ddd, *J* = 13.8, 9.5, 6.7 Hz, 1H), 2.04 – 1.71 (m, 7H), 1.69 – 1.49 (m, 6H), 1.48 – 1.23 (m, 6H), 1.12 (dd, *J* = 12.2, 5.8 Hz, 1H), 1.02 (d, *J* = 6.5 Hz, 3H), 1.00 – 0.93 (m, 1H), 0.92 (s, 3H), 0.71 (s, 3H). $^{13}$C NMR (126 MHz, DMSO) δ 173.82, 172.61, 71.02, 70.44, 66.25, 54.92, 48.60, 46.15, 45.74, 41.53, 41.37, 35.32, 35.20, 35.05, 34.90, 34.70, 34.40, 32.50, 31.70, 30.41, 28.57, 27.31, 26.22, 22.82, 22.63, 17.12, 12.35. HR-MS (ESI-TOF-MS): $[M-H]^-$ calcd. for $C_{28}H_{45}N_2O_7^-$ 478.3174; found 478.3171.

**Lithocholoyl taurine (Tau-LCA)**

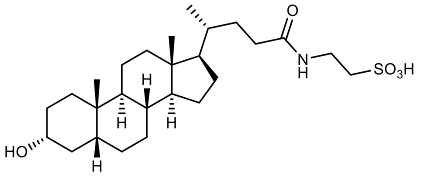


To a 100 mL round-bottom flask charged with a stir bar was added THF (25 mL) followed by lithocholic acid (750 mg, 1.99 mmol, 1.00 mmol). The resulting solution was cooled to 0 °C in an ice/water bath, and triethylamine (0.33 mL, 2.39 mmol, 1.2 equiv.) and ethyl chloroformate (0.23 mL, 2.40 mmol, 1.21 equiv.) were added dropwise. The reaction was then allowed to warm to room temperature for 1.5 h, after which the white slurry was cooled to 0 °C. A solution of taurine (383 mg, 3.07 mmol, 1.54 equiv) and sodium bicarbonate (251 mg, 2.99 mmol, 1.50 equiv.) in water (25 mL) was then added dropwise *via* cannula. The solution was allowed to warm to room temperature with stirring for 2 h. The reaction was concentrated under reduced pressure and acidified with 2 M HCl. The resulting white solid was collected, washed with water, and recrystallized from hot EtOH/EtOAc (2:3)[7] as a white powder (250 mg, 26% yield).

Analytical data: $^1$H NMR (500 MHz, MeOD) δ 3.61 (t, *J* = 6.9 Hz, 2H), 3.53 (d, *J* = 4.9 Hz, 1H), 2.97 (t, *J* = 6.8 Hz, 2H), 2.37 – 2.24 (m, 1H), 2.19 – 2.08 (m, 1H), 2.06 – 1.98 (m, 1H), 1.90 (tt, *J* = 9.4, 4.2 Hz, 2H), 1.79 (ddd, *J* = 17.1, 12.3, 7.6 Hz, 3H), 1.69 – 1.55 (m, 2H), 1.52 – 1.23 (m, 12H), 1.23 – 1.04 (m, 5H), 1.04 – 0.98 (m, 1H), 0.98 – 0.92 (m, 6H), 0.69 (s, 3H). $^{13}$C NMR (126 MHz, DMSO) δ 172.08, 69.88, 56.08, 55.54, 50.61, 42.27, 41.54, 36.31, 35.43, 35.39, 35.17, 34.93, 34.22, 32.53, 31.44, 30.40, 27.72, 26.90, 26.17, 23.86, 23.29, 20.41, 18.30, 11.88. HR-MS (ESI-TOF-MS): $[M-H]^-$ calcd. for $C_{26}H_{44}NO_5S^-$ 482.2945; found 482.2940.

### Supplemental Tables

**Table S1.** Plasmids used in this study.

| Plasmid | Protein Sequence Accession | Description | Source |
|---|---|---|---|
| pET21b_Cperfringens_cBSH | P54965 (UniProt) | Cperfringens_cBSH - FLAG - His$_6$ | Parasar *et al* 2019[8] |
| pET21b_Tfaecis_mBSH | WP_161832809.1 (NCBI) | Tfaecis_mBSH - FLAG - His$_6$ | This study |
| pET21b_GCA-900066495 sp902362365_mBSH | MGYG000186436_00677 (UHGG) | GCA-900066495 sp902362365_mBSH - FLAG - His$_6$ | This study |
| pET21b_Rtimonensis_mBSH | MGYG000252833_00592 (UHGG) | Rtimonensis_mBSH - FLAG - His$_6$ | This study |
| pET21b_Cbaratii_mBSH | MGYG000278967_00137 (UHGG) | Cbaratii_mBSH - FLAG - His$_6$ | This study |
| pET21b_Cdifficile_mBSH | MGYG000269495_01068 (UHGG) | Cdifficile_mBSH - FLAG - His$_6$ | This study |
| pET21b_Heffluvii_mBSH | MGYG000203535_02775 (UHGG) | Heffluvii_mBSH - FLAG - His$_6$ | This study |
| pET21b_UBA7185 sp900055645_mBSH | MGYG000000285_02149 (UHGG) | UBA7185 sp900055645_mBSH - FLAG - His$_6$ | This study |
| pET21b_Bmassiliensis_mBSH_wt | MGYG000001487_00309 (UHGG) | Bmassiliensis_mBSH_wt - FLAG - His$_6$ | This study |
| pET21b_Bmassiliensis_mBSH_S144A | | Bmassiliensis_mBSH_S144A - FLAG - His$_6$ | This study |
| pET21b_Bmassiliensis_mBSH_Q290A | | Bmassiliensis_mBSH_Q290A - FLAG - His$_6$ | This study |
| pET21b_Bmassiliensis_mBSH_N337A | | Bmassiliensis_mBSH_N337A - FLAG - His$_6$ | This study |
| pET21b_Bmassiliensis_mBSH_S144A_Q290A | | Bmassiliensis_mBSH_S144A_Q290A - FLAG - His$_6$ | This study |
| pET21b_Bmassiliensis_mBSH_S144A_N337A | | Bmassiliensis_mBSH_S144A_N337A - FLAG - His$_6$ | This study |
| pET21b_Bmassiliensis_mBSH_Q290A_N337A | | Bmassiliensis_mBSH_Q290A_N337A - FLAG - His$_6$ | This study |
| pET21b_Bmassiliensis_mBSH_S144A_Q290A_N337A | Derived from pEТ21b_Bmassiliensis_mBSH_wt | Bmassiliensis_mBSH_S144A_Q290A_N337A - FLAG - His$_6$ | This study |
| pET21b_GCA-900066495 sp902362365_cBSH | MGYG000186436_00318 (UHGG) | GCA-900066495 sp902362365_cBSH - His$_6$ | This study |
| pET21b_Bwadsworthia_mBSH | MGYG000043481_01420 (UHGG) | Bwadsworthia_mBSH - His$_6$ | This study |

**Table S2.** Primers used in this study.

| Primer | Sequence (5' to 3') |
|---|---|
| Tf_mBSH_fwd | CTAGCATATGCGTAAGCTGTACTTCAACG |
| Tf_mBSH_rev | CTAGCTCGAGCTTATCGTCGTCATCCTTGTAATCCTTCTTG TAGATGGTGTTACC |
| GCA-900066495 sp902362365_mBSH_fwd | CTAGCATATGATGAGGAAAATTTACTTCAACGGCAAC |
| GCA-900066495 sp902362365_mBSH_rev | CTAGCTCGAGCTTATCGTCGTCATCCTTGTAATCTTTTTA TAGATCGTGTTGCCCTCC |
| GCA-900066495 sp902362365_mBSH_internalSequencing | CCTTTGGGTATGTAGGCACCAGC |
| Rtimonensis_mBSH_fwd | CTAGCATATGATGACCAAGAAACTTTATTTTAACGG |
| Rtimonensis_mBSH_rev | CTAGCTCGAGCTTATCGTCGTCATCCTTGTAATCGGCTTTA AAGATTGTGTTGCCCTC |
| Rtimonensis_mBSH_internalSequencing | CCTCATGAAAGATGAGGGTA |
| Cbaratii_mBSH_fwd | CTAGCATATGATGACCAAGAAACTTTATTTTAACGG |
| Cbaratii_mBSH_rev | CTAGCTCGAGCTTATCGTCGTCATCCTTGTAATCGGC TTTAAAGATTGTGTTGCCCTC |
| Cbaratii_mBSH_internalSequencing | CGTAAAAGAATACTTAAA |
| Cdifficile_mBSH_fwd | CTAGCATATGATGAAAGAAAAAATTTATTACAATGG CA |
| Cdifficile_mBSH_rev | CTAGCTCGAGCTTATCGTCGTCATCCTTGTAATCCTT CAGAGAATACAGTACCTCG |
| Cdifficile_mBSH_internalSequencing | CGATAACAACCGTGAGTT |
| Heffluvii_mBSH_fwd | CTAGCATATGATGGATAAAATCTTATACCATGGTACC |
| Heffluvii_mBSH_rev | CTAGCTCGAGCTTATCGTCGTCATCCTTGTAATCTCG TTTGTAAATCGTCTCACCG |
| Heffluvii_mBSH_internalSequencing | CGGAACTGGCCGGTCAGTTTCTG |
| UBA7185 sp900055645_mBSH_fwd | CTAGCATATGATGGGTAAGACGTTGTACTTGAATGG |
| UBA7185 sp900055645_mBSH_rev | CTAGCTCGAGCTTATCGTCGTCATCCTTGTAATCCTC TGCGTTATAAACAACGGTGC |
| UBA7185 sp900055645_mBSH_internalSequencing | CGAGAATCATGTCAAACTGG |
| Bmassiliensis_mBSH_fwd | CTAGCATATGATGCATACGATTTATTATAACGGGA |
| Bmassiliensis_mBSH_rev | CTAGCTCGAGCTTATCGTCGTCATCCTTGTAATCTTT GTTGCGTTTAAAAATGGTTT |
| Bmassiliensis_mBSH_internalSequencing | CGCTACCACCACCGCTTGAAA |
| Bmassiliensis_mBSH_S144A_mutagenic_fwd | GTTCTTATTCATGCCGCCGGCCACATGGGCA |
| Bmassiliensis_mBSH_S144A_mutagenic_rev | TGCCCATGTGGCCGGCGGCATGAATAAGAAC |
| Bmassiliensis_mBSH_Q290A_mutagenic_fwd | AGATGGTTCGCCGGCGGGCCGAACGGCC |
| Bmassiliensis_mBSH_Q290A_mutagenic_rev | GGCCGTTCGGCCCGCCGGCGAACCATCT |
| Bmassiliensis_mBSH_N337A_mutagenic_fwd | CGCGTCTCCGGCACAATGCGCCAGTAGCTGC |
| Bmassiliensis_mBSH_N337A_mutagenic_rev | GCAGCTACTGGCGCATTGTGCCGGAGACGCG |
| GCA-900066495 sp902362365_cBSH_fwd | CTAGCATATGATGTGTACCGCGCTAACCTTAAC |
| GCA-900066495 sp902362365_cBSH_rev | CTAGCTCGAGGTTTTCACTGTTAAACACCTGGGTATC |
| Bwadsworthia_mBSH_fwd | CTAGCATATGATGGGCAAAGCAACGCAGAT |
| Bwadsworthia_mBSH_rev | CTAGCTCGAGTTCCAGCGACCCGTATACTAATTC |
| Bwadsworthia_mBSH_internalSequencing | AAAGCAAATTATACCAAGG |

**Table S3.** Human gut bacterial homologs of *Turicibacter faecis* mBSH found in the uhgp-100. (Excel file)

**Table S4.** Kinetic parameters for characterized BSHs. Parameters were fit according to the Michaelis—Menten equation. Values are reported as mean ± sem of *n* = 3 independent experiments.

| Enzyme | Substrate | $k_{cat}$ ($min^{-1}$) | $K_m$ ($\mu M$) | $k_{cat}/K_m$ ($min^{-1}\mu M^{-1}$) |
|---|---|---|---|---|
| *Turicibacter faecis* TC023 mBSH[a] | TCA | 176.4 ± 5.0 | 49.2 ± 2.1 | 3.6 ± 0.2 |
| | TCDCA | 819.4 ± 29.1 | 88.2 ± 5.9 | 9.3 ± 0.4 |
| | TDCA | 1001.8 ± 47.3 | 129.8 ± 14.1 | 7.8 ± 0.5 |
| GCA-900066495 sp902362365 mBSH[a] | TCA | 94.4 ± 4.8 | 27.0 ± 3.9 | 3.8 ± 0.2 |
| | TCDCA | 453.8 ± 1.1 | 83.2 ± 7.1 | 5.7 ± 0.4 |
| | TDCA | 957.3 ± 80.2 | 157.7 ± 24.9 | 6.1 ± 0.4 |
| GCA-900066495 sp902362365 cBSH[b] | TCA | 648.7 ± 23.0 | 353.0 ± 42.9 | 1.9 ± 0.2 |
| | TCDCA | 930.7 ± 26.6 | 942.4 ± 101.1 | 1.0 ± 0.1 |
| | TDCA | 1223.2 ± 63.5 | 858.6 ± 123.8 | 1.5 ± 0.2 |

[a]Determined using 30 nM BSH with reaction monitoring by LC-MS.

[b]Determined using 200 nM BSH with reaction monitoring by ninhydrin assay.

**Table S5.** BLAST queries for taurine catabolism and sulfidogenesis proteins. (Excel file)

**Table S6.** Taurine catabolism and sulfidogenesis proteins identified in genomes harboring *Turicibacter faecis* mBSH homologs. (Excel file)

**Table S7.** cBSH sequences from genomes harboring *Turicibacter faecis* mBSH homologs. (Excel file)

**Table S8.** BLAST queries used for amidohydrolase 3 domain network annotation. (Excel file)

**Table S9.** Catalytic residue-filtered mBSH protein sequences from validated clusters of amidohydrolase 3 domain network. (Excel file)

**Table S10.** Catalytic residue-filtered cBSH protein sequences from uhgp-90. (Excel file)

**Table S11.** Widespread, differentially abundant *mbsh* genes associated with cardiovascular disease (CVD).

| Gene | Taxonomy | Cardiovascular disease association |
|---|---|---|
| MGYG000010551_01338 | *Medterranibacter gnavus* | Positive |
| MGYG000008858_01273 | *Medterranibacter gnavus* | Positive |
| MGYG000233073_01757 | *Ruthenibacterium lactatiformans* | Positive |
| MGYG000071840_02437 | *Enterocloster bolteae* | Positive |
| MGYG000071704_02575 | *Hungatella sp005845265* | Positive |
| MGYG000113645_01364 | *Bilophila wadsworthia* | Negative |
| MGYG000092686_02218 | *Bilophila wadsworthia* | Negative |
| MGYG000152805_01493 | *Bilophila wadsworthia* | Negative |
| MGYG000051378_02177 | *Bilophila wadsworthia* | Negative |

**Table S12.** Taurine catabolism proteins identified in the genomes containing positively associated CVD *mbsh* genes. (Excel file)

### Supplementary Figures

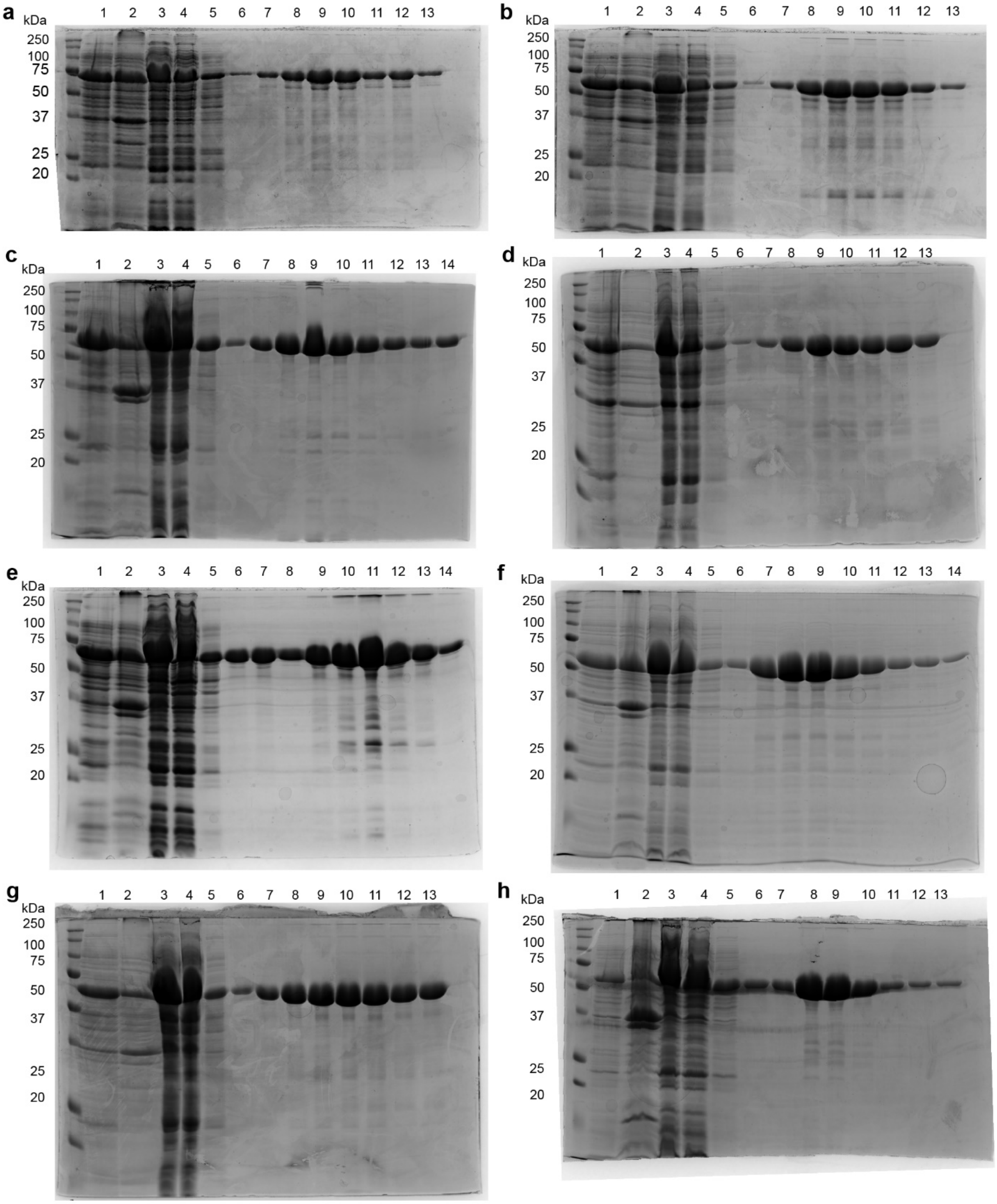


1: Total lysate 2: Insoluble fraction 3: Soluble fraction 4: Flow-through 5: 20 mM imidazole wash 6: 50 mM imidazole wash
7: 100 mM imidazole elution 8: 150 mM imidazole elution 9: 200 mM imidazole elution 10-14: 250 mM imidazole elution

**Figure S1.** Overexpression and purification of mBSH proteins from (a) *T. faecis*, (b) *Clostridium baratii*, (c) *Romboutsia timonensis*, (d) *Clostridioides difficile*, (e) *GCA-900066495 sp902362365*, (f) *Hungatella effluvii,* (g) *UBA7185 sp900055645,* and (h) *Beduini massiliensis*.

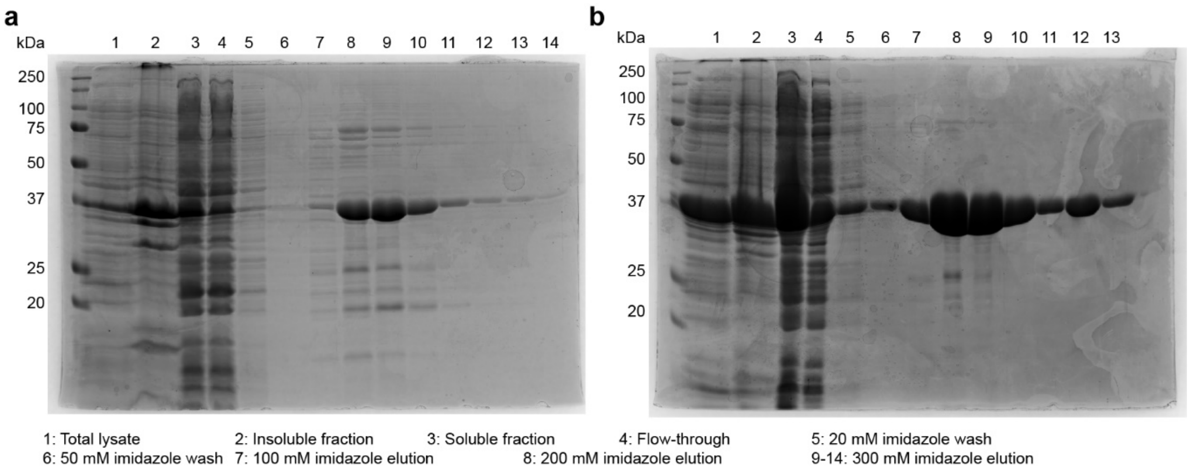


**Figure S2.** Overexpression and purification of cysteine hydrolase BSHs from (a) *Clostridium perfringens* and (b) *GCA-900066495 sp902362365*.

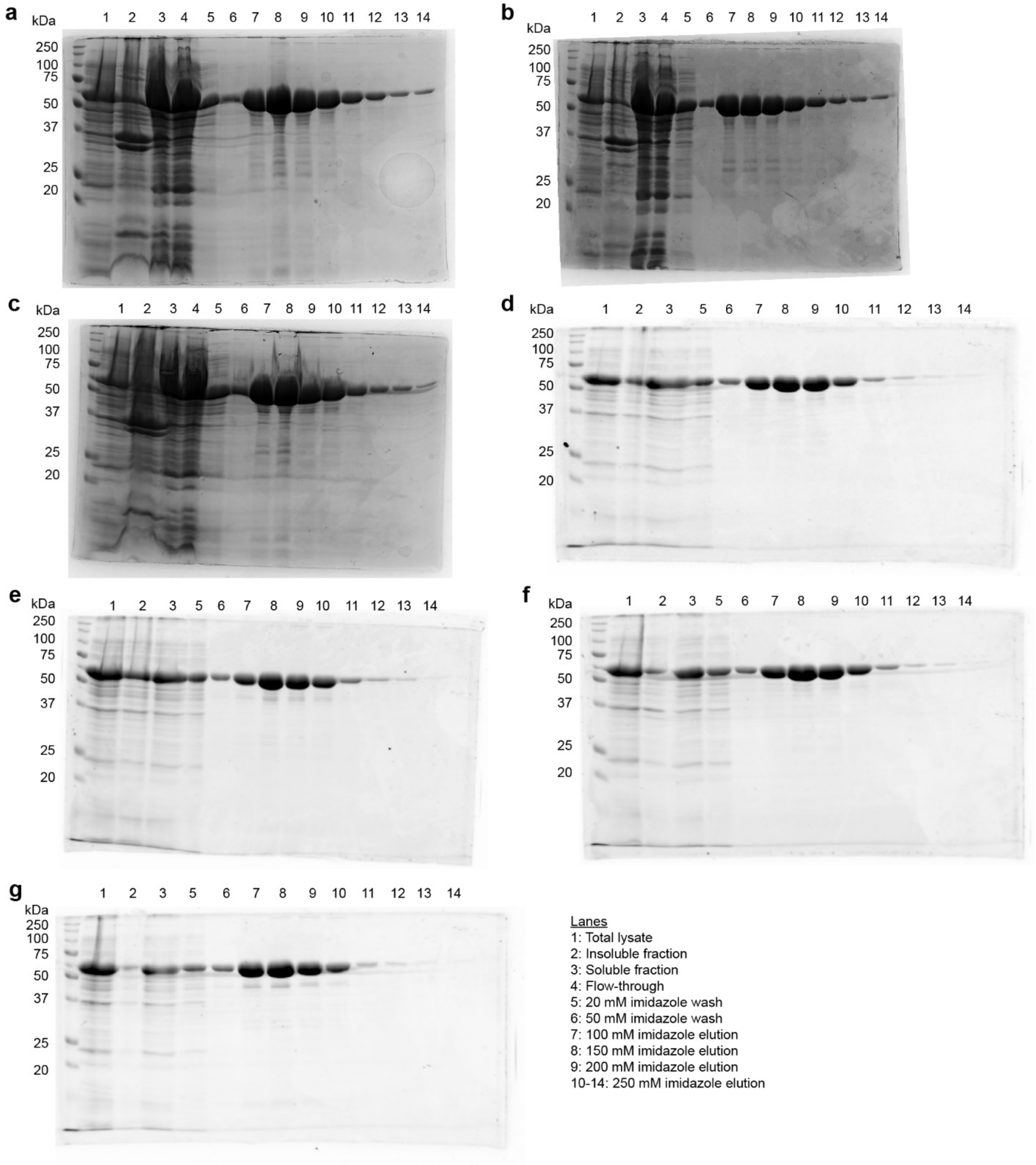


**Figure S3.** Overexpression and purification of *Beduini massiliensis* mutant mBSH variants (a) S144A; (b) Q290A; (c) N337A; (d) S144A, Q290A; (e) S144A, N337A; (f) Q290A, N337A; and (f) S144A, Q290A, N337A.

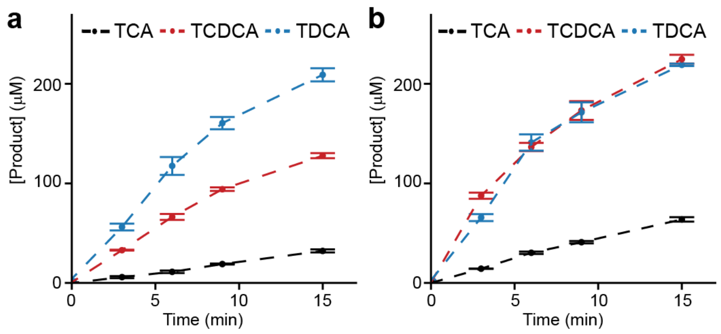


**Figure S4.** Determination of initial rate conditions for (a) *GCA-900066495 sp902362365* and (b) *Turicibacter faecis* mBSH in $NaP_i$ (25 mM, 1% DMSO, pH = 6.8) with BSH (30 nM) and taurine-conjugated bile acid (300 µM). Data representative of *n* = 3 independent experiments. All values are reported as mean ± sem.

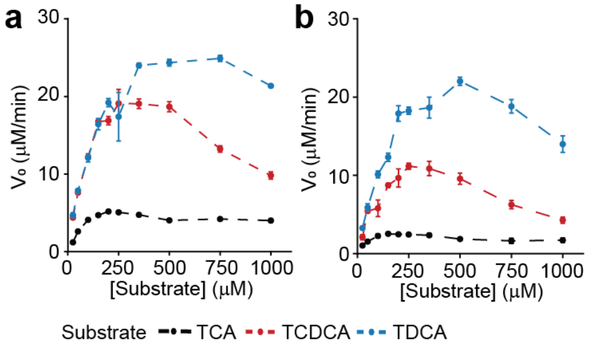


**Figure S5.** Extended range of kinetic for measurements indicating substrate inhibition for (a) *Turicibacter faecis and* (b) *GCA-900066495* mBSH in $NaP_i$ (25 mM, 1% DMSO, pH = 6.8) with BSH (30 nM) and taurine-conjugated bile acid (0 - 1000 µM). Data representative of *n* = 3 independent experiments. All values are reported as mean ± sem.

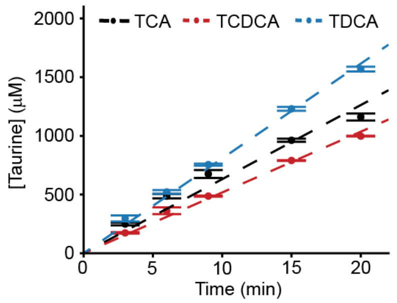


**Figure S6.** Determination of initial rate conditions for *GCA-900066495 sp902362365* cBSH in $NaP_i$ (25 mM, % 1 DMSO, pH = 6.8) with BSH (200 nM) and taurine-conjugated bile acid (2 mM). Data representative of *n* = 3 independent experiments. All values are reported as mean ± sem.

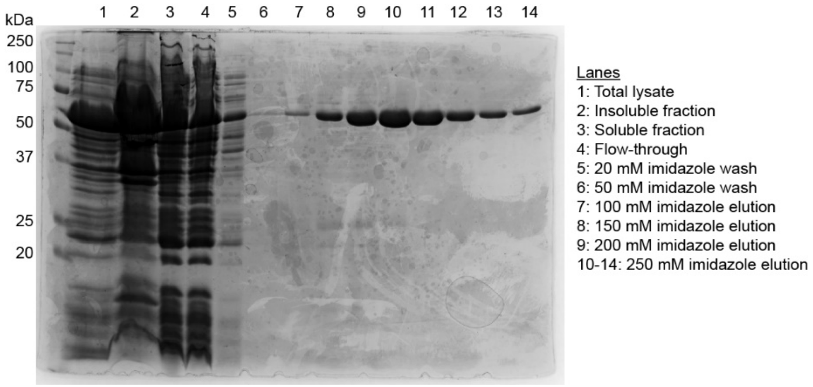


**Figure S7.** Overexpression and purification of *Bilophila wadsworthia* mBSH.

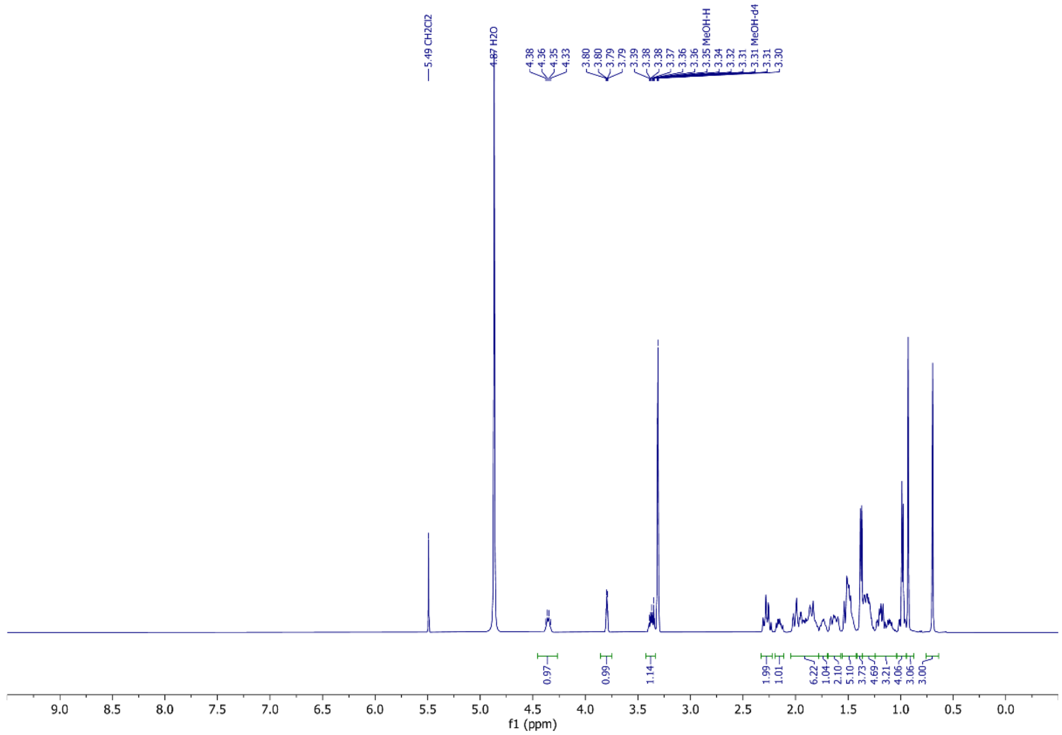


**Figure S8.** [1]H NMR spectrum of Ala-CDCA (MeOH-*d*4, 500 MHz)

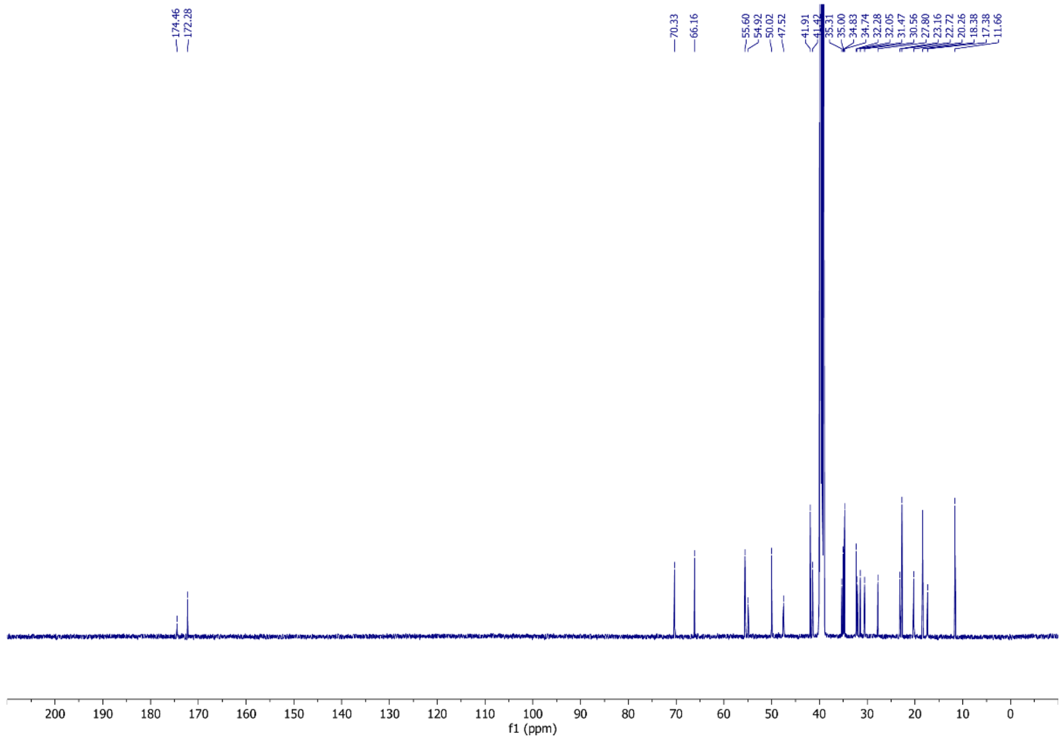


**Figure S9.** $^{13}C$ NMR of Ala-CDCA (DMSO-*d*6, 126 MHz)

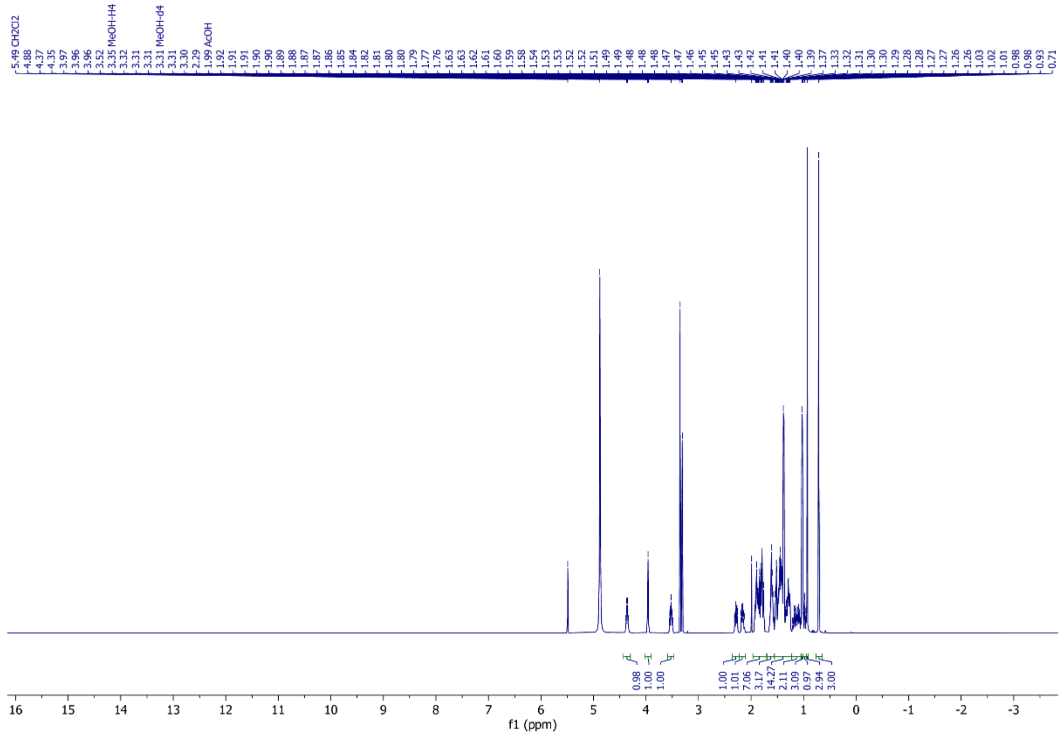


**Figure S10.** $^{1}H$ NMR spectrum of Ala-DCA (MeOH-*d*4, 500 MHz)

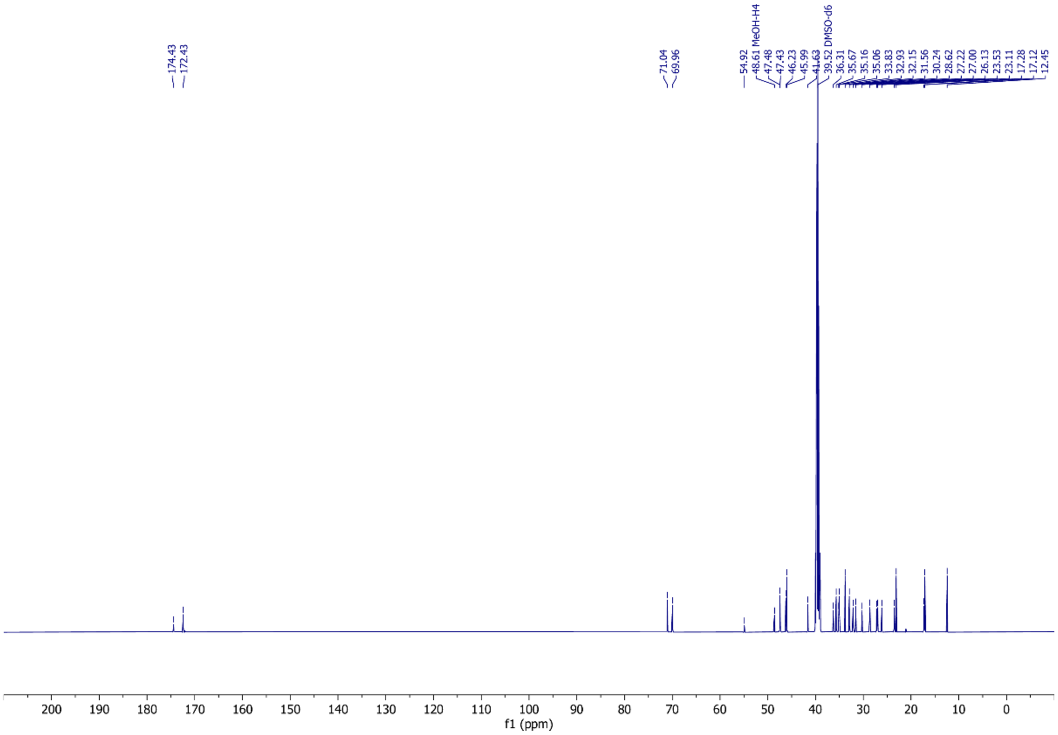


**Figure S11.** $^{13}$C NMR of Ala-DCA (DMSO-*d*6, 126 MHz)

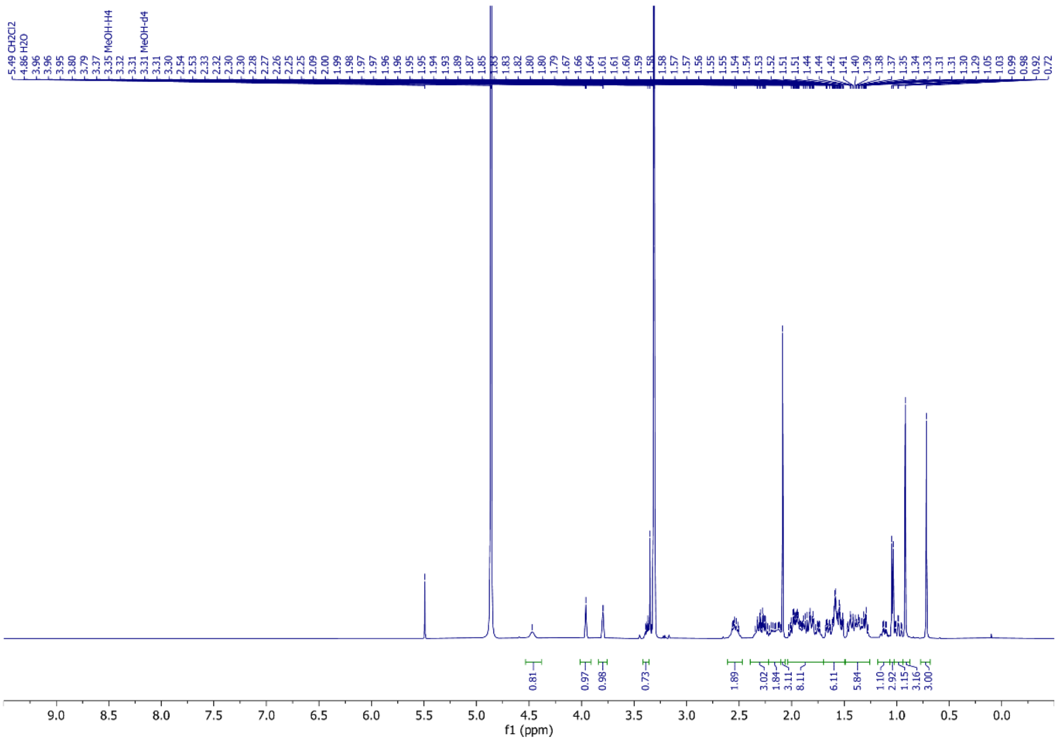


**Figure S12.** $^{1}$H NMR spectrum of Met-CA (MeOH-*d*4, 500 MHz)

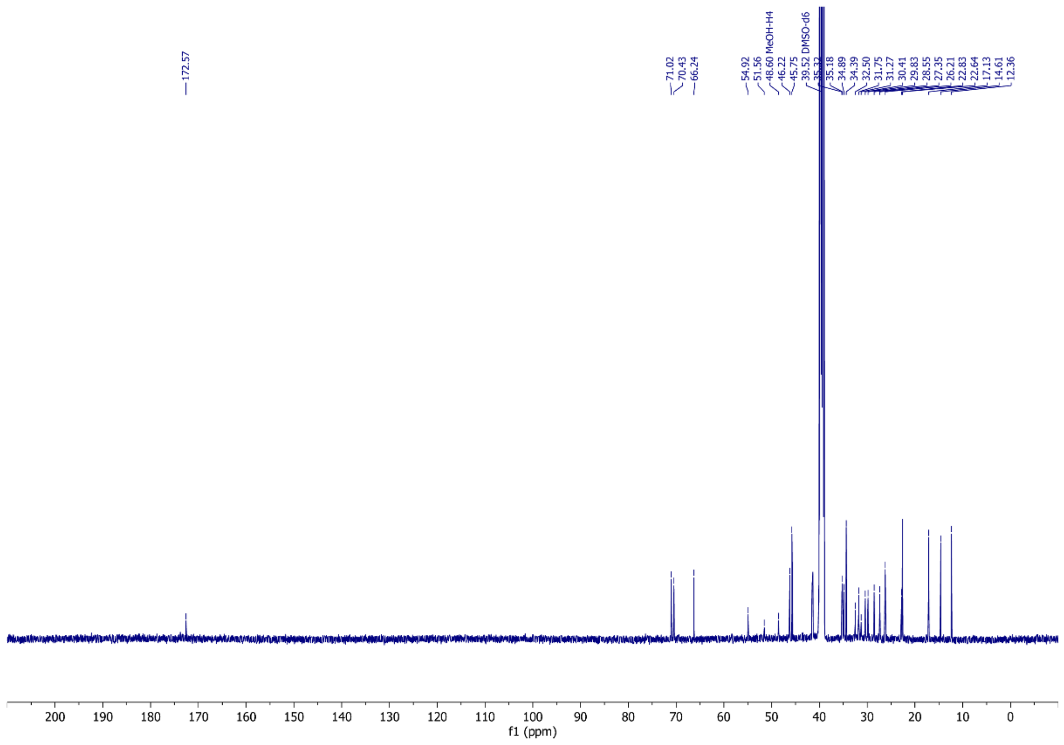

**Figure S13.** $^{13}C$ NMR of Met-CA (DMSO-*d*6, 126 MHz)

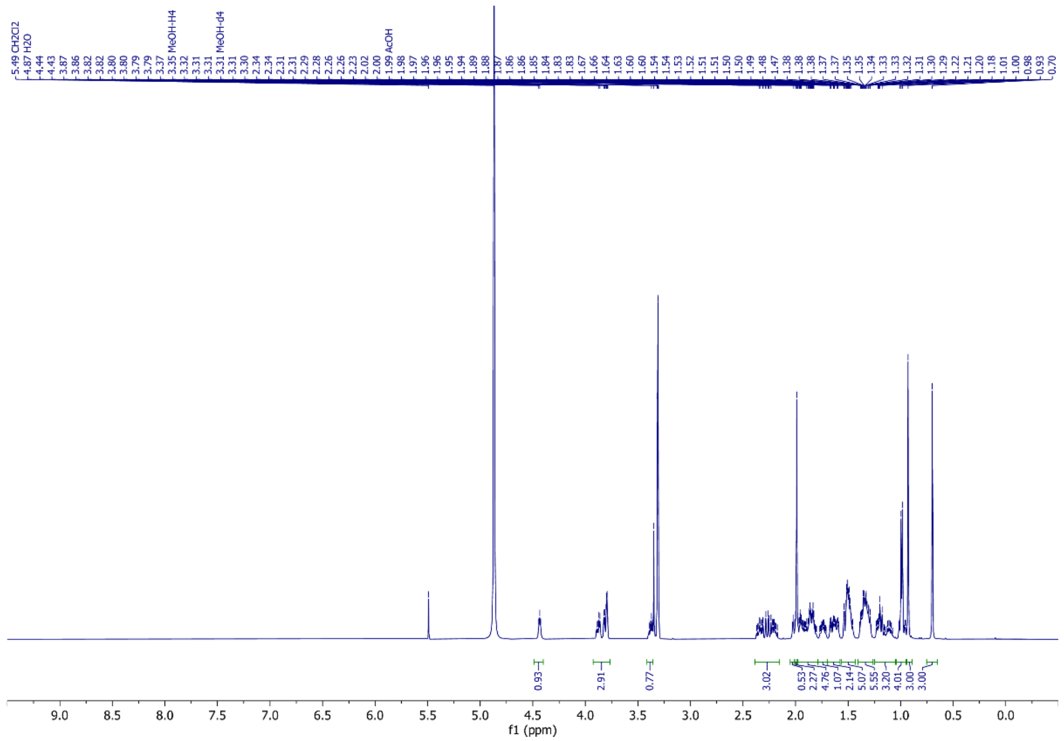

**Figure S14.** $^{1}H$ NMR spectrum of Ser-CDCA (MeOH-*d*4, 500 MHz)

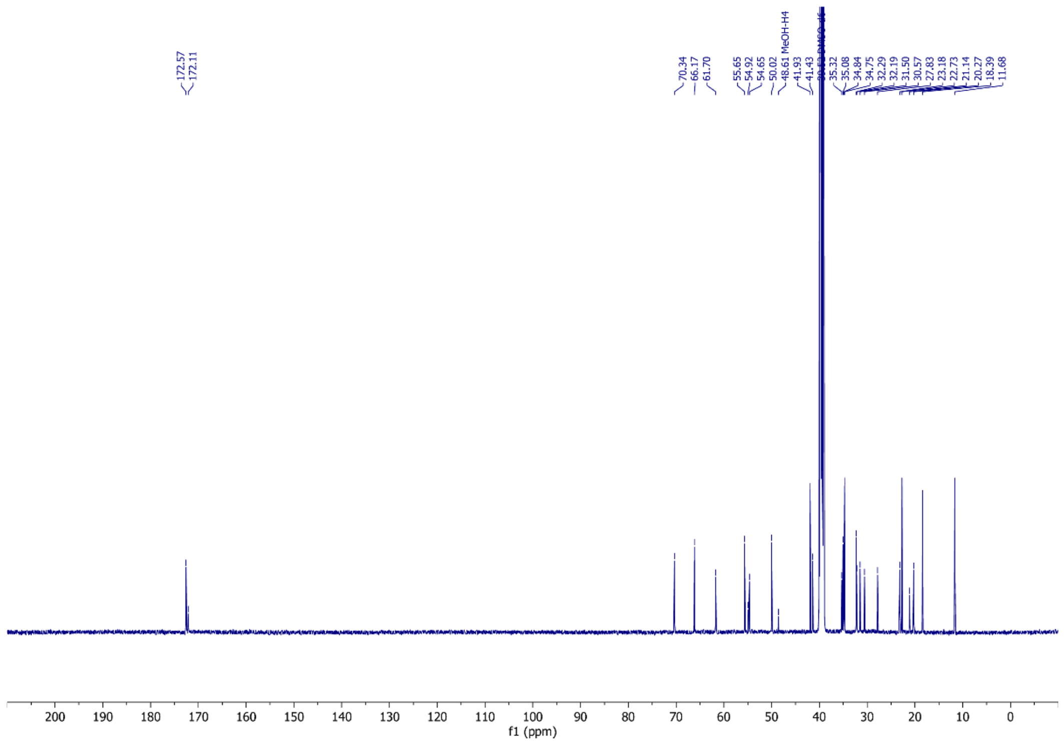

**Figure S15.** $^{13}C$ NMR of Ser-CDCA (DMSO-*d*6, 126 MHz)

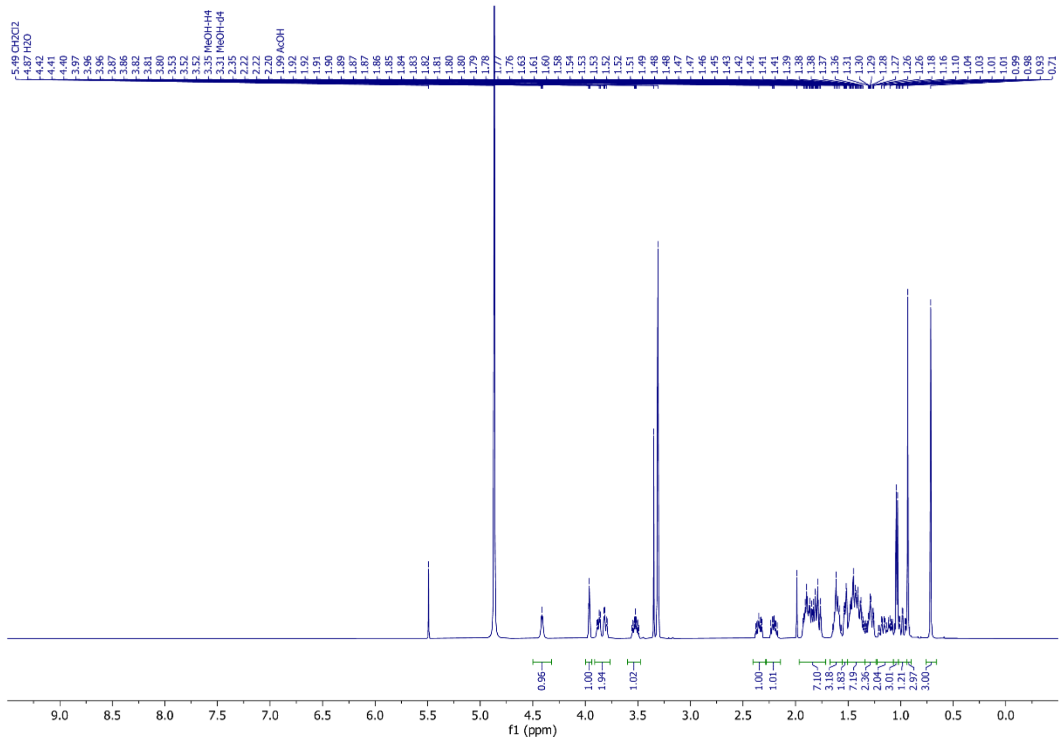

**Figure S16.** $^{1}H$ NMR spectrum of Ser-DCA (MeOH-*d*4, 500 MHz)

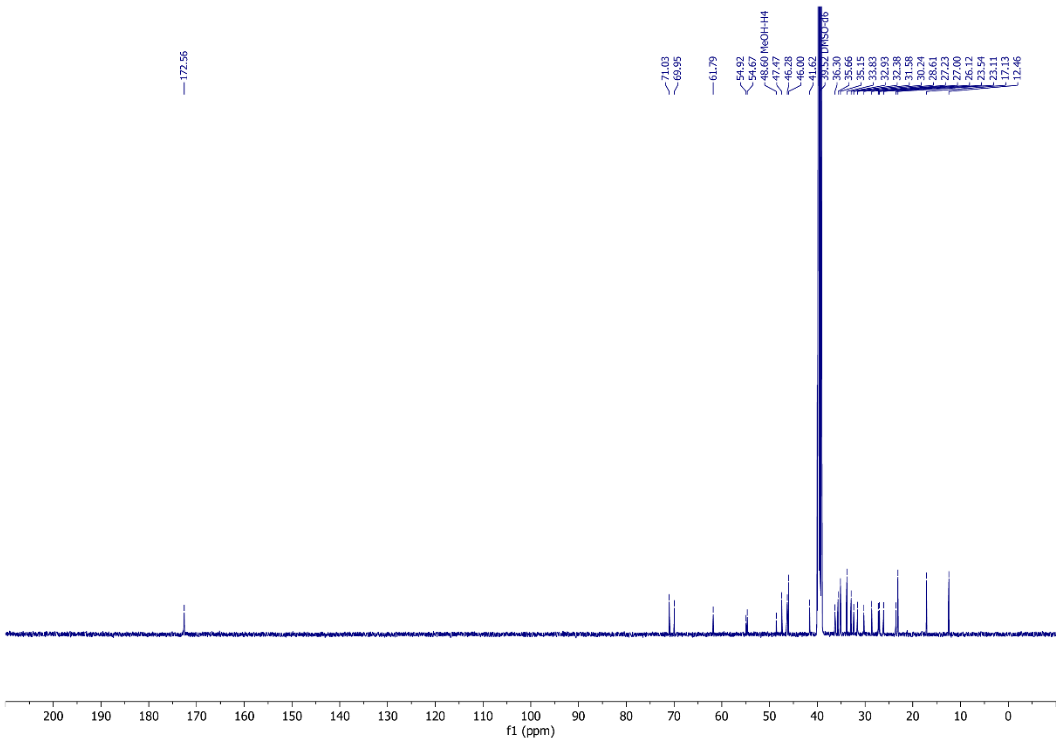


**Figure S17.** $^{13}$C NMR of Ser-DCA (DMSO-*d*6, 126 MHz)

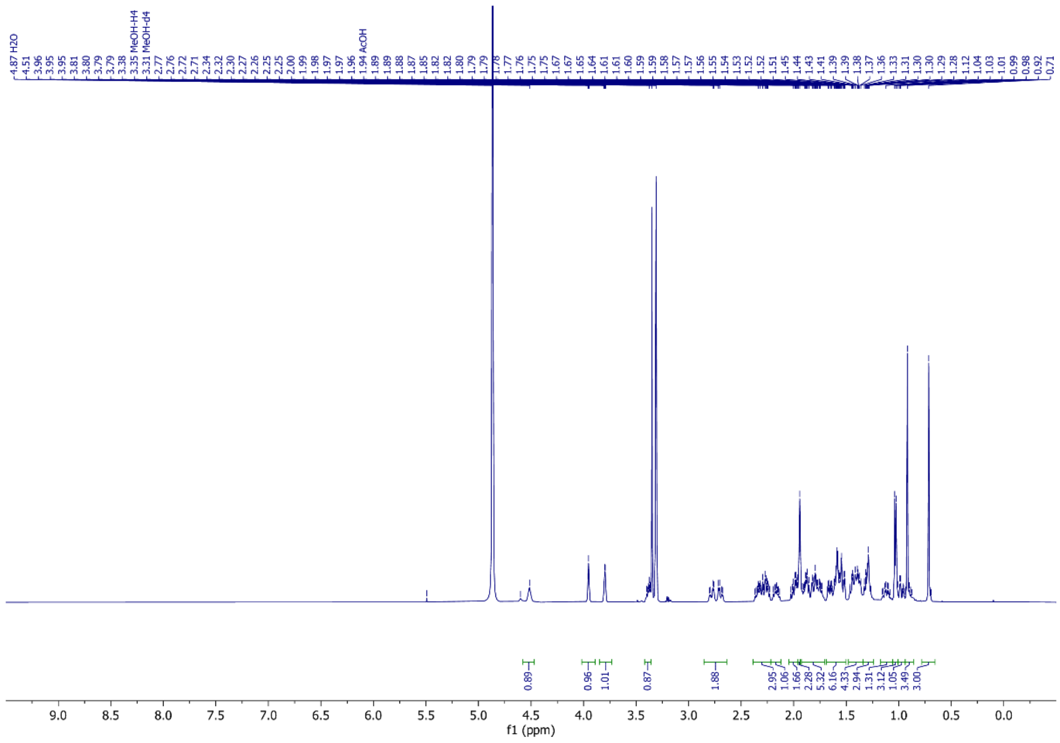


**Figure S18.** $^{1}$H NMR spectrum of Asn-CA (MeOH-*d*4, 500 MHz)

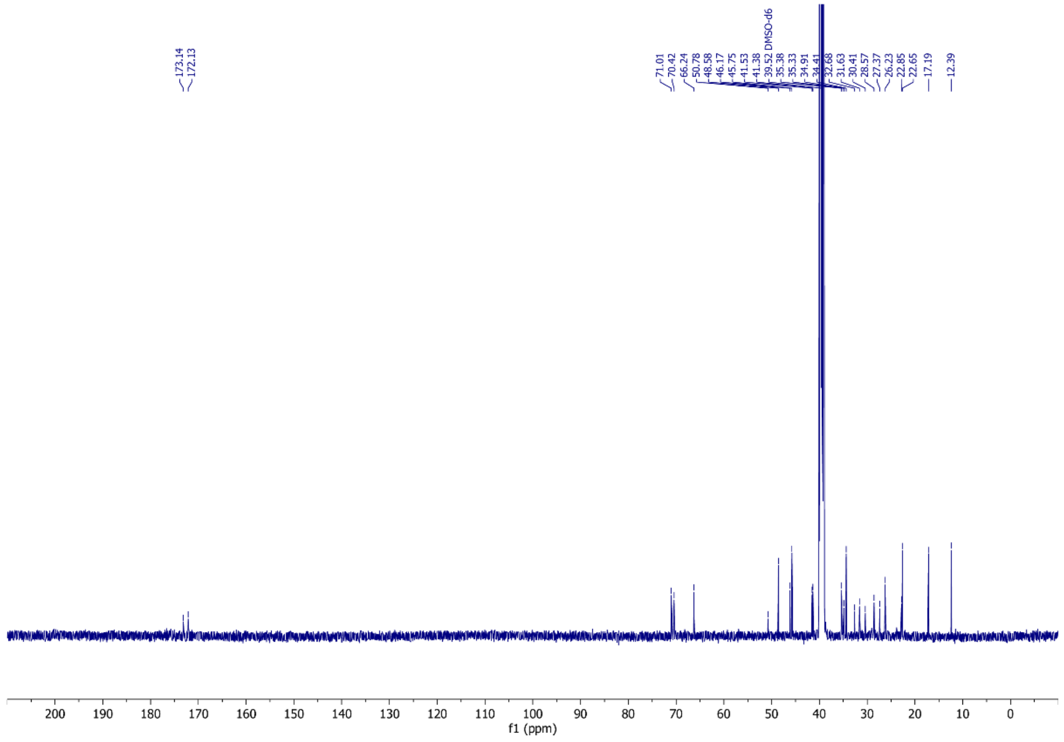


**Figure S19.** $^{13}$C NMR of Asn-CA (DMSO-*d*6, 126 MHz)

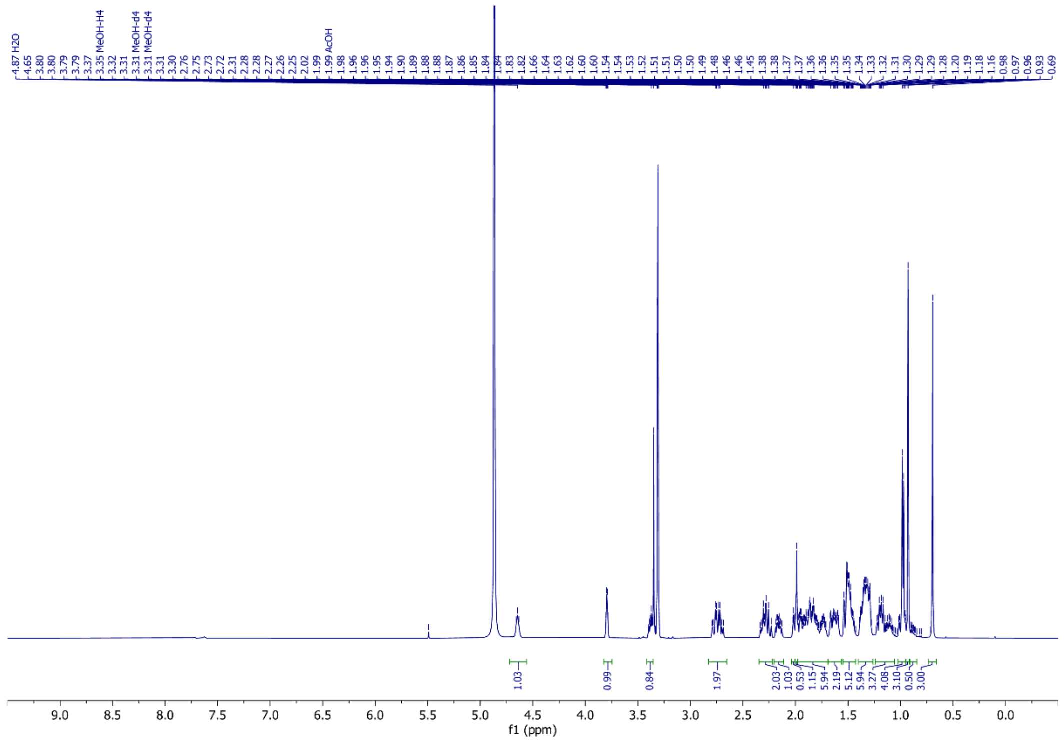


**Figure S20.** $^{1}$H NMR spectrum of Asn-CDCA (MeOH-*d*4, 500 MHz)

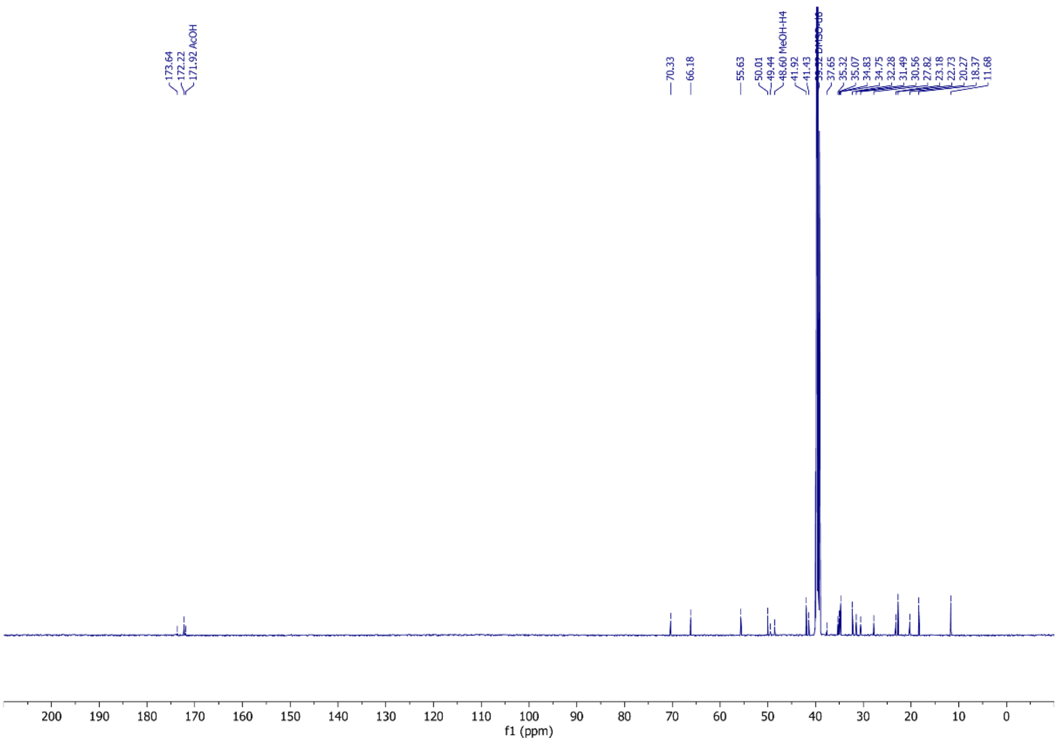


**Figure S21.** $^{13}$C NMR of Asn-CDCA (DMSO-*d*6, 126 MHz)

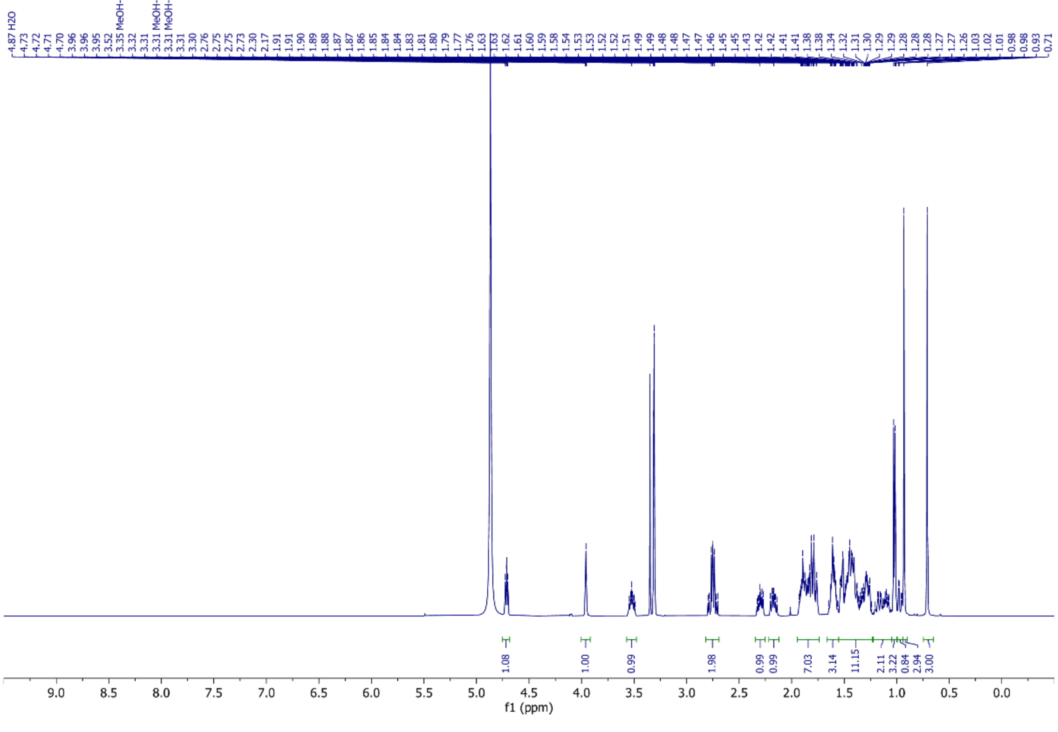


**Figure S22.** $^{1}$H NMR spectrum of Asn-DCA (MeOH-*d*4, 500 MHz)

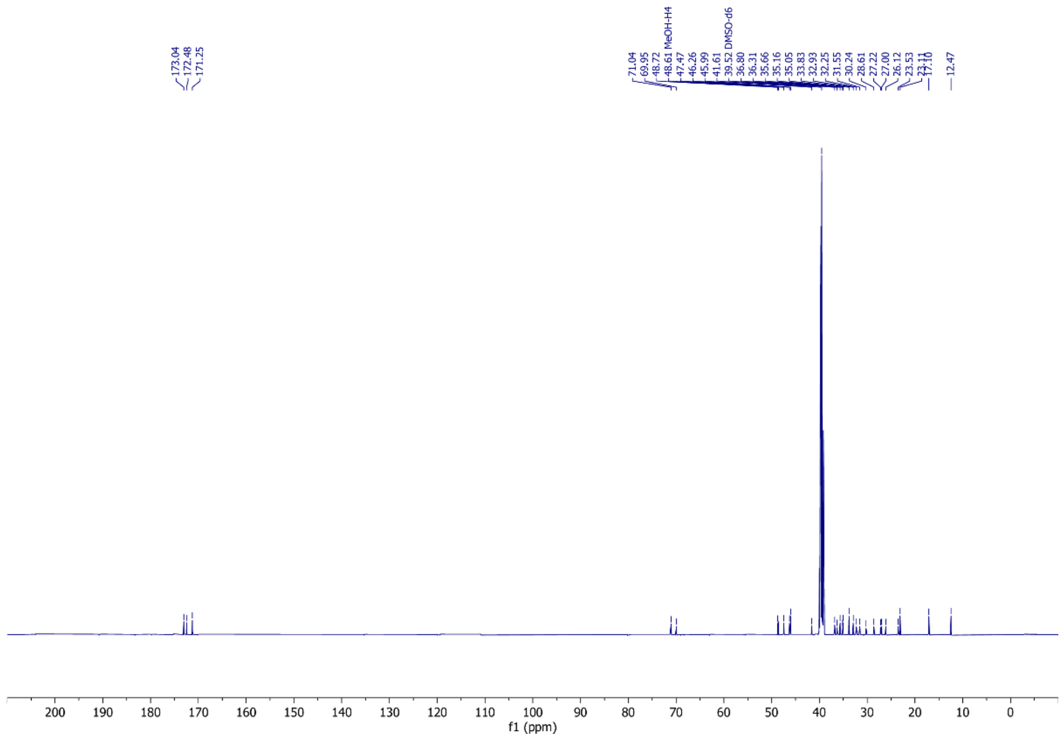


**Figure S23.** $^{13}C$ NMR of Asn-DCA (DMSO-*d*6, 126 MHz)

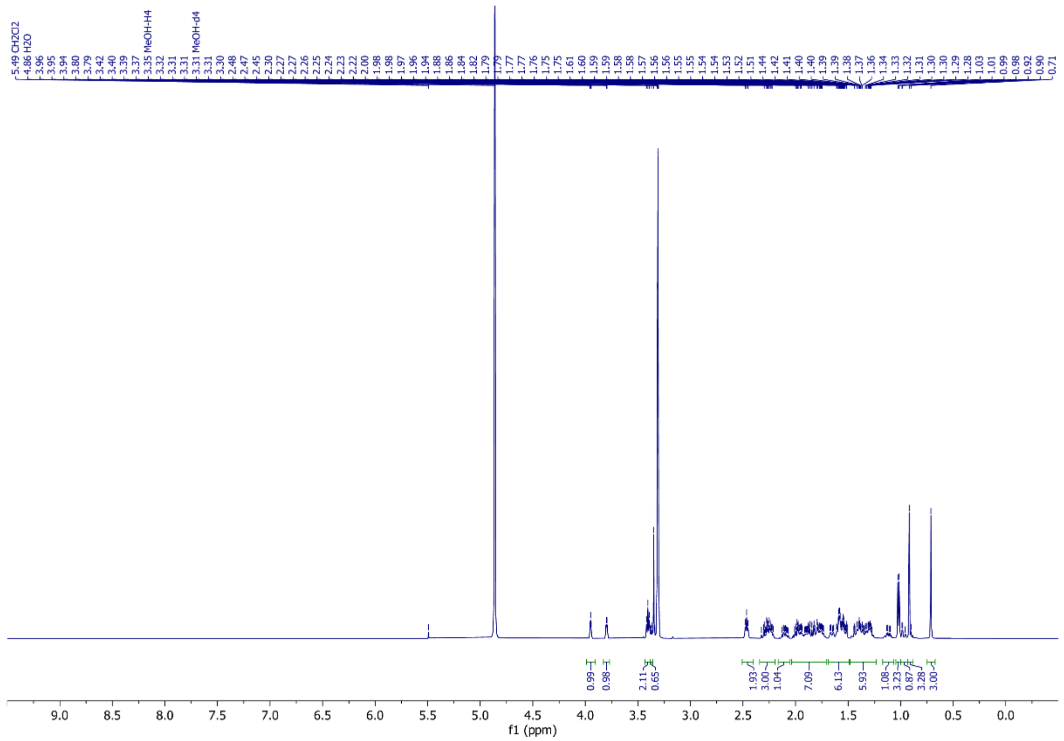


**Figure S24.** $^{1}H$ NMR spectrum of β-Ala-CA (MeOH-*d*4, 500 MHz)

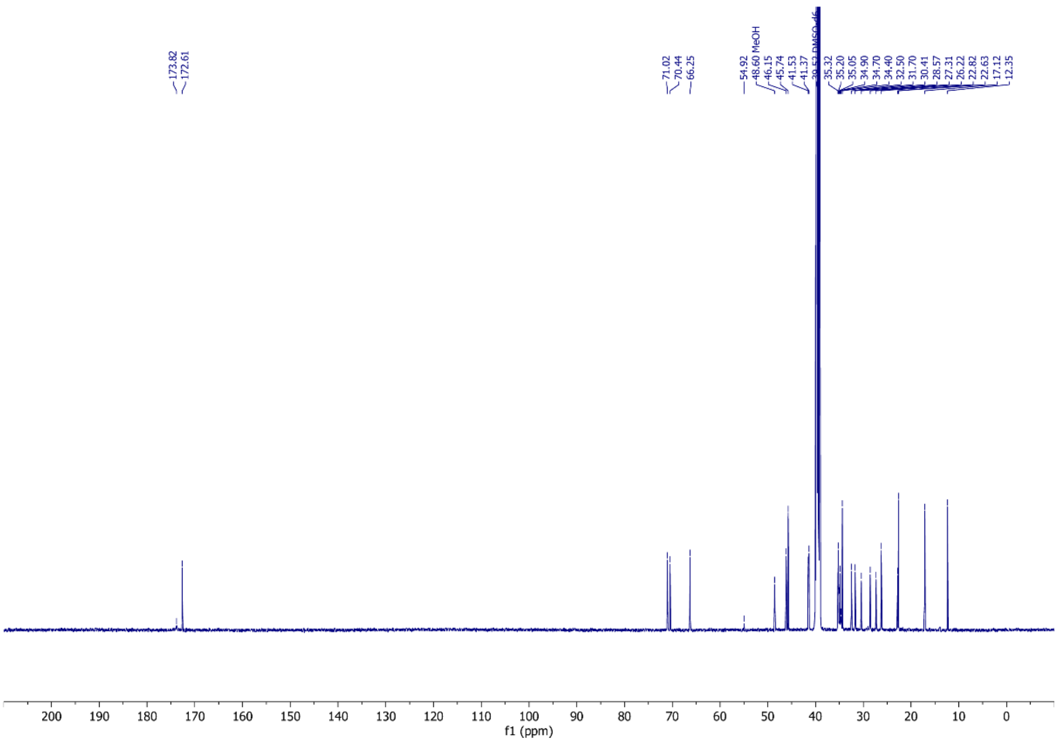


**Figure S25.** $^{13}C$ NMR of β-Ala-CA (DMSO-*d*6, 126 MHz)

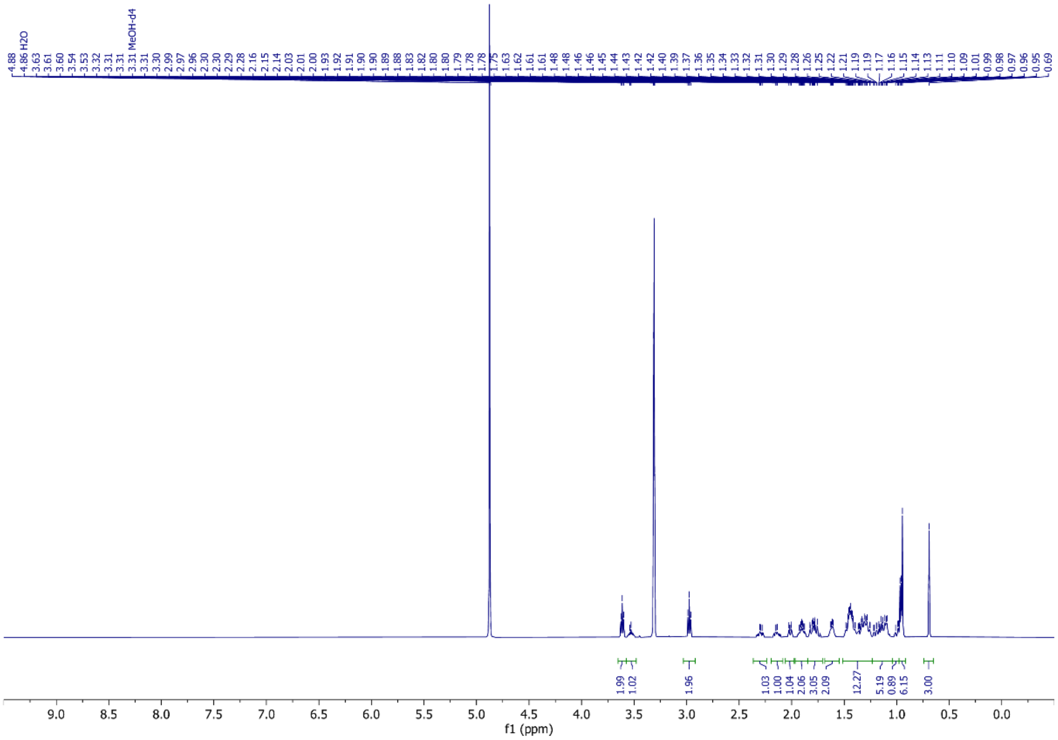


**Figure S26.** $^{1}H$ NMR spectrum of Tau-LCA (MeOH-*d*4, 500 MHz)

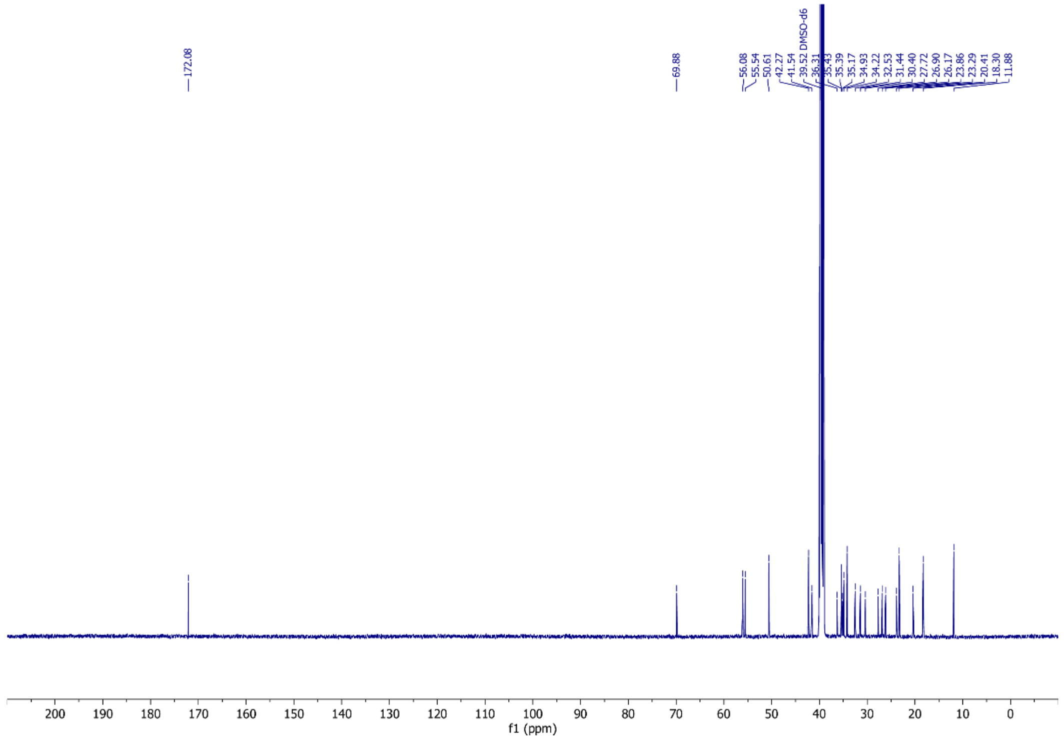


**Figure S27.** $^{13}$C NMR of Tau-LCA (DMSO-*d*6, 126 MHz)
